# DR5 disulfide bonding as a sensor and effector of protein folding stress

**DOI:** 10.1101/2024.03.04.583390

**Authors:** Mary E. Law, Zaafir M. Dulloo, Samantha R. Eggleston, Gregory P. Takacs, Grace M. Alexandrow, Mengxiong Wang, Hanyu Su, Bianca Forsyth, Chi-Wu Chiang, Abhisheak Sharma, Siva Rama Raju Kanumuri, Olga A. Guryanova, Jeffrey K. Harrison, Boaz Tirosh, Ronald K. Castellano, Brian K. Law

**Affiliations:** Department of Pharmacology & Therapeutics, University of Florida, 32610; Department of Chemistry, University of Florida, Gainesville, FL, 32611; Department of Pharmaceutics, University of Florida, Gainesville, FL, 32610; UF Health Cancer Center, University of Florida, Gainesville, FL, 32610; Department of Radiation Biology, Stanford University, Stanford, CA, 94305; Institute of Molecular Medicine, College of Medicine, National Cheng Kung University, Tainan, Taiwan; Department of Biochemistry, Case Western Reserve University, Cleveland, OH, 44106

**Author notes:** Equally contributing co-first authors (Mary E. Law, Zaafir M. Dulloo).

## Abstract

New agents are needed that selectively kill cancer cells without harming normal tissues. The TRAIL ligand and its receptors, DR5 and DR4, exhibit cancer-selective toxicity, but TRAIL analogs or agonistic antibodies targeting these receptors have not received FDA approval for cancer therapy. Small molecules for activating DR5 or DR4 independently of protein ligands may bypass some of the pharmacological limitations of these protein drugs. Previously described Disulfide bond Disrupting Agents (DDAs) activate DR5 by altering its disulfide bonding through inhibition of the Protein Disulfide Isomerases (PDIs) ERp44, AGR2, and PDIA1. Work presented here extends these findings by showing that disruption of single DR5 disulfide bonds causes high-level DR5 expression, disulfide-mediated clustering, and activation of Caspase 8-Caspase 3 mediated pro-apoptotic signaling. Recognition of the extracellular domain of DR5 by various antibodies is strongly influenced by the pattern of DR5 disulfide bonding, which has important implications for the use of agonistic DR5 antibodies for cancer therapy. Disulfide-defective DR5 mutants do not activate the ER stress response or stimulate autophagy, indicating that these DDA-mediated responses are separable from DR5 activation and pro-apoptotic signaling. Importantly, other ER stressors, including Thapsigargin and Tunicamycin also alter DR5 disulfide bonding in various cancer cell lines and in some instances, DR5 mis-disulfide bonding is potentiated by overriding the Integrated Stress Response (ISR) with inhibitors of the PERK kinase or the ISR inhibitor ISRIB. These observations indicate that the pattern of DR5 disulfide bonding functions as a sensor of ER stress and serves as an effector of proteotoxic stress by driving extrinsic apoptosis independently of extracellular ligands.

## Introduction

Cancer remains one of the most lethal diseases, making the identification of safer and more effective therapies urgent. Identification of cancer drug targets that are essential for malignant cells, but not normal cells, is key. Targeting proteins involved in the folding and maturation of oncoproteins, but not “house-keeping” proteins, holds great promise. Protein Disulfide Isomerases (PDIs) comprise a family of 22 human enzymes that play essential roles in the folding of secreted and membrane proteins [1]. Previous work showed that PDIs may be favorable targets for anticancer agents [2–6]. However, much of this work focused on canonical PDIs with CXXC active site motifs and little is known about non-canonical PDIs that possess CXXS active site trapping motifs that lack the second, resolving cysteine. Previous work indicated that bicyclic thiosulfonate compounds termed Disulfide bond Disrupting Agents (DDA) bind to the PDIs PDIA1, ERp44, AGR2, and AGR3 through their active site Cys residues [7]. DDAs block client binding to PDIA1 and ERp44 and prevent disulfide-mediated AGR2 dimerization. Further, mutation of the active site Cys residues of ERp44 and AGR2 ablate binding to biotinylated DDAs. Collectively, these results suggest that DDAs inhibit the catalytic activity of PDIA1, ERp44, AGR2, and AGR3 by covalently modifying their active site Cys residues.

Importantly, DDAs show significant activity against breast tumors and metastatic lesions in animal models without affecting surrounding stromal cells or normal tissues [8, 9]. Tumor cell death occurred through apoptosis, and DDA-mediated apoptosis was associated with downregulation of the HER-family oncoproteins EGFR, HER2, and HER3 and upregulation and activation of DR5, a receptor for the pro-apoptotic ligand TRAIL. However, significant questions remain regarding DDA modes of anticancer action, determinants of cancer responsiveness to DDAs, and the features controlling DDA safety and metabolic stability. The work presented here was designed to address these questions. The results reveal that DR5 plays a central role in the cancer-selective, pro-apoptotic effects of the DDAs, that DR5 levels and signaling activity through the Caspase 8-Caspase 3 axis are controlled by the state of DR5 disulfide bonding, and that multiple inducers of endoplasmic reticulum protein folding stress alter DR5 disulfide bonding. These observations suggest that DR5 functions as both a sensor and effector of proper disulfide bond formation in proteostasis.

## Results

### DDA-triggered selective ER retention (sERr) is associated with elevated DR5 levels and signaling

The DDAs used herein are presented in Fig. 1A. We proposed that DDAs exhibit rapidly reversible covalent bonding to protein thiols by disulfide bond formation, with the exception of the target PDIs that form stable disulfide bonds with DDAs [10]. In further support of this premise, we incubated T47D cell extracts with the biotinylated DDA probe Bio-Pyr-DTDO alone or combined with a 100-fold excess of the unlabeled DDA competitors shown in Fig. 1B. Bands recognized by Bio-Pyr-DTDO were identified as PDIA1, ERp44, and AGR2 by mass spectrometry and immunoblot as reported previously [7]. Endogenous biotinylated proteins are observed in the absence of Bio-Pyr-DTDO treatment (asterisks). As expected, Bio-Pyr-DTDO binding was blocked by the more reactive, less selective DDAs DTDO, D5DO, D7DO, and RBF3. In contrast, the less reactive, more selective DDA tcyDTDO did not affect Bio-Pyr-DTDO binding, nor did the thiol-reactive deubiquitinase inhibitor b-AP15 [11]. The thiol-reactive compound *N*-ethylmaleimide prevented Bio-Pyr-DTDO binding to DDA targets. These observations support the selectivity of bicyclic DDAs against a subset of PDIs.

**Fig. 1:**
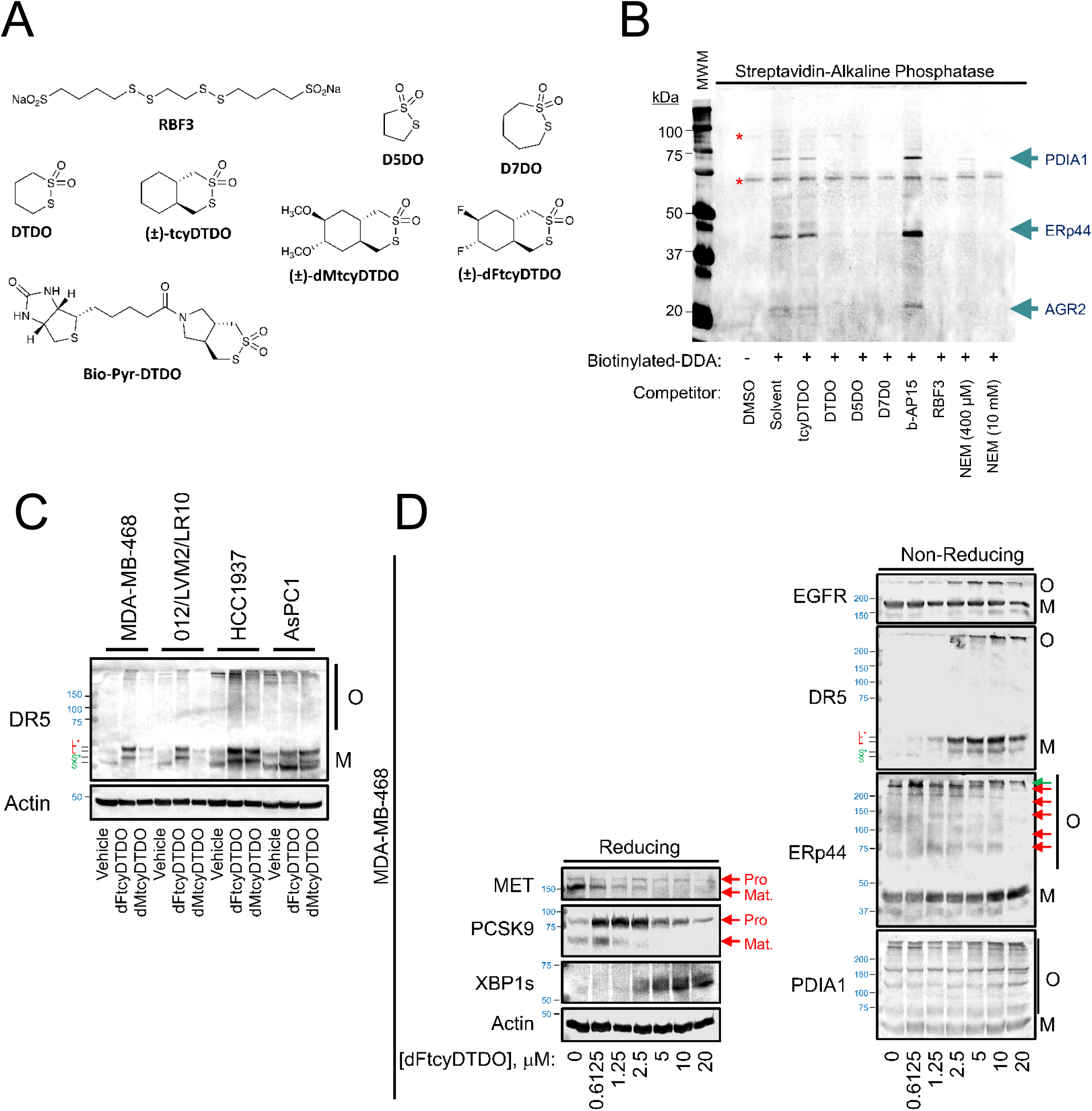
DDA compounds that selectively inhibit AGR2, PDIA1, and ERp44 block the maturation of select transmembrane and secreted proteins, but strongly upregulate DR5. A. Chemical structures of representative DDAs used in the manuscript. B. Demonstration of the selectivity of biotinylated DDA binding to the target proteins PDIA1, ERp44, and AGR2. Extracts from T47D cells were incubated with the indicated competitors for 2h and then incubated for 1 h with biotinylated-DDA, followed by sample analysis by gel electrophoresis and blotting with Streptavidin-Alkaline Phosphatase detection. C. Non-reducing immunoblot analysis of the effect of 24 h treatment of the indicated cells with the specified DDAs at 2.5 μM each. M represents monomeric DR5 isoforms and O represents disulfide-bonded DR5 oligomeric complexes. S and L refer to the short and long forms of DR5 and S′ and L′ refer to the same DR5 isoforms with altered electrophoretic mobility caused by DDA treatment. Actin serves as a loading control. D. Left panel, reducing immunoblot analysis of MDA-MB-468 cells treated with increasing dFtcyDTDO concentrations showing higher expression of XBP1s and decreased levels of the mature forms and increased relative levels of the pro-forms of MET and PCSK9. Right panel, non-reducing immunoblot analysis using the indicated antibodies. Red arrows represent oligomeric ERp44 isoforms lost upon dFtcyDTDO treatment and the green arrow represents high molecular mass ERp44 isoforms elevated by dFtcyDTDO treatment. O and M represent the Oligomeric and Monomeric protein isoforms in panels C and D.

Also consistent with previous work, the bicyclic DDAs dFtcyDTDO and dMtcyDTDO increased the levels of DR5, and immunoblot analysis under non-reducing conditions showed an electrophoretic mobility shift of monomeric DR5 and an increase in disulfide bonded oligomeric forms of DR5 (Fig. 1C). A previous study from the Tirosh laboratory showed that under Endoplasmic Reticulum (ER) stress conditions, trafficking of some mis-disulfide bonded transmembrane receptor tyrosine kinases became arrested in the ER through the formation of large disulfide bonded complexes involving ERp44 [12]. This mechanism was termed selective ER retention, or sERr. Since DDAs induce ER stress and inhibit ERp44 client binding [7], we examined whether DDA treatment activates or inhibits sERr. Increasing dFtcyDTDO blocked maturation of MET, an established marker of sERr and also prevented maturation of PCSK9 through its auto-cleavage (Fig. 1D). This was associated with increased expression of the ER stress marker XBP1s. Analysis of the same samples under non-reducing conditions showed that increasing dFtcyDTDO concentrations increased EGFR oligomerization, and elevated levels of monomeric and oligomeric DR5. dFtcyDTDO had a modest effect on PDIA1 client binding. In contrast, dFtcyDTDO blocked the formation of lower molecular mass ERp44 disulfide bonded complexes with clients (red arrows), while very high mass ERp44 complexes (green arrow) were elevated. This observation suggests that DDAs caused sERr. However, the observation that high levels of mis-disulfide bonded monomeric DR5 accumulated suggests that DR5 may evade sERr.

The DDA dMtcyDTDO also induced sERr as indicated by near complete blockade of MET maturation (Fig. 2A). This effect was not altered by signaling inhibitors of mTORC1 (rapamycin), Akt (MK2206), EGFR (Gefitinib), or EGFR and HER2 (Lapatinib). The PERK kinase suppresses protein synthesis under ER stress conditions in part through phosphorylation of eIF2α [12], and previous work [12] showed that sERr is strongly potentiated by PERK inhibition. In MDA-MB-468 cells, Thapsigargin induced a partial block of MET processing that was strongly potentiated by the PERK inhibitor GSK2606414 (hereafter called PERKi) as expected. Since DDAs activate sERr, and sERr is potentiated by PERKi, cell viability studies were performed to examine the effect of DDA/PERKi combination treatment. While PERKi alone had little effect on cell viability, PERKi strongly potentiated dFtcyDTDO cytotoxicity in MDA-MB-468 breast cancer cells (2B, left panel) and WM793 melanoma cells (2B, right panel). sERr was initially investigated primarily in HepG2 hepatoma cells [12], so we compared the combinatorial effects of dMtcyDTDO and PERKi in HepG2 and MDA-MB-468 cells. In both lines PERKi alone had no effect on MET or PCSK9 processing (Fig. 2C). However, PERKi increased the levels of unprocessed MET in both lines and elevated levels of unprocessed PCSK9 in HepG2 cells. This is consistent with PERKi permitting continued synthesis of nascent MET and PCSK9 under ER stress conditions. Combinatorial DDA/PERKi treatment was associated with increased Caspase 8 cleavage, which could explain the enhanced toxicity to cancer cells. PERKi did not strongly potentiate DDA upregulation of DR5, but potentiated Caspase 8 cleavage/activation (Fig. 2D). DR5 knockout partially blunted Caspase 8 cleavage. The compound ISRIB [13] negates the integrated stress response (ISR) by overcoming the effects of eIF2α phosphorylation [14]. When combined with dMtcyDTDO, ISRIB and PERKi produced similar enhancements in the levels of unprocessed MET and PCSK9 (Fig. 2E). This is consistent with both agents overcoming ISR triggered by DDA treatment. Combinatorial DDA/PERKi activation of Caspase 8 in MDA-MB-468 cells was not altered by forced CDCP1 expression, which disrupts cell-cell adhesion and confers suspension growth [15], (Supplemental Fig. S1A), and PERKi actions were not mimicked by inhibition of eEF2K [6] that controls translation initiation (Supplemental Fig. S1B). Together, these observations suggest that PERKi enhances DDA toxicity to cancer cells by potentiating DR5 pro-apoptotic signaling rather than upregulating DR5 expression.

**Fig. 2:**
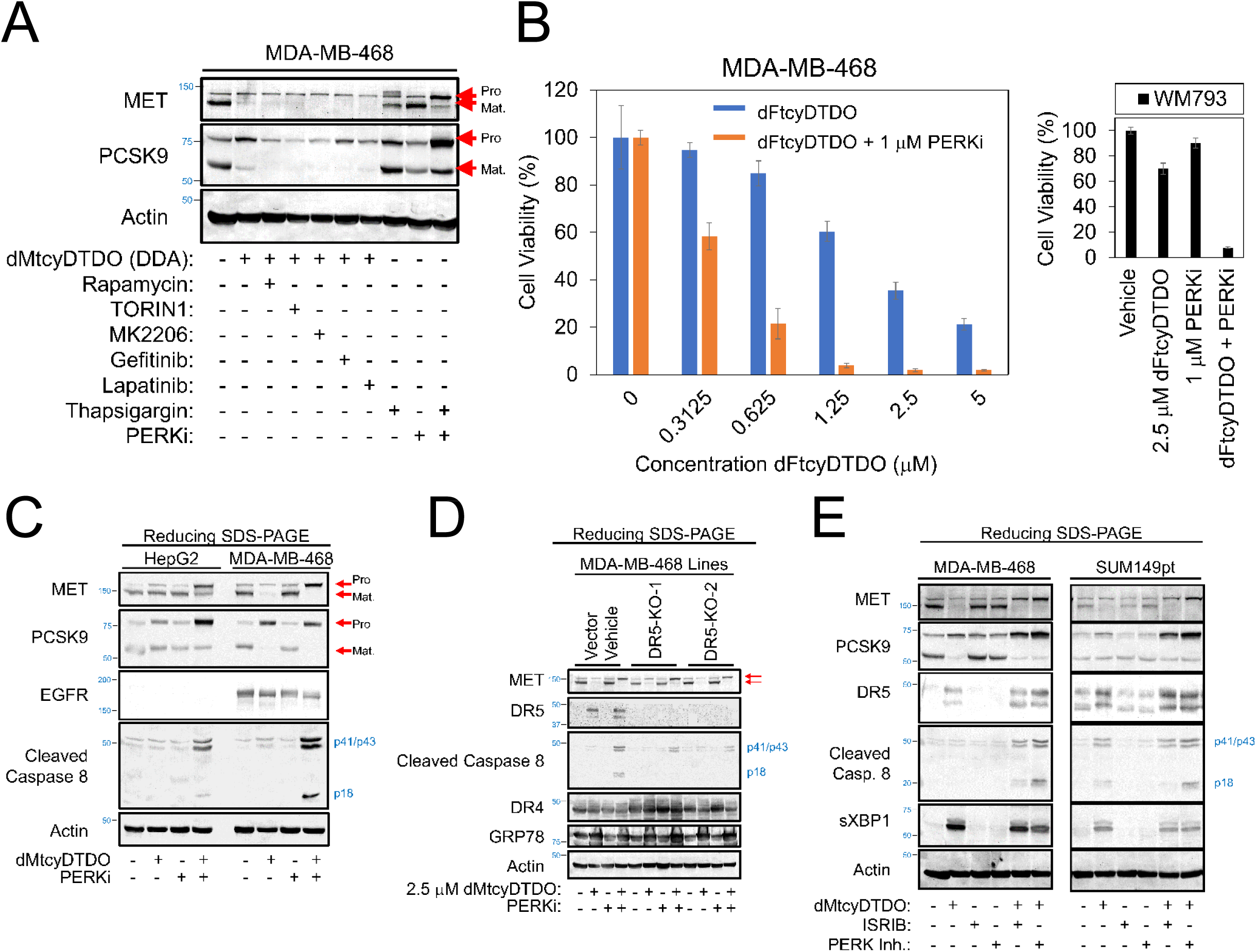
PERK inhibition amplifies the pro-apoptotic effects of DDAs on cancer cell lines. A. Reducing immunoblot analysis of MDA-MB-468 cells treated for 24 h with the indicated combinations of dMtcyDTDO (2.5 μM), Rapamycin (100 nM), TORIN1 (100 nM), MK2206 (5 μM), Gefitinib (10 μM), Lapatinib (10 μM), Thapsigargin (400 nM), and PERKi (1μM). Red arrows denote pro- or mature protein isoforms. B. MTT cell viability assays of MDA-MB-468 cells (left panel) or WM793 cells (right panel) treated for 72 h as indicated. Data are plotted as the average (N = 6), with error bars representing standard deviation. C. Reducing immunoblot analysis of HepG2 or MDA-MB-468 cells treated for 24 h as indicated with 2.5 μM dMtcyDTDO or 1 μM PERKi. Red arrows denote pro- or mature protein isoforms. D. Reducing immunoblot analysis of the indicated cell lines treated as specified for 24 h with dMtcyDTDO (2.5 μM) or PERKi (1 μM). E. Reducing immunoblot analysis of MDA-MB-468 cells or SUM149pt cells treated for 24 h as indicated with dMtcyDTDO (2.5 μM), ISRIB (200 nM), or PERKi (1μM).

### Multiple ER stress inducers alter DR5 disulfide bonding

Since ERp44 is a DDA target, we examined if ER stress alters DR5 disulfide bonding in the absence of ERp44, or in HepG2 ERp44 knockout cells in which wild type or catalytically null (C29S) versions of ERp44 were reintroduced. The most notable effect was that the ER stressor Thapsigargin and PERKi had little effect alone, but irrespective of the presence or absence of ERp44, Thapsigargin + PERKi decreased DR5 electrophoretic mobility and increased its levels (Fig. 3A). This suggests that while DDAs are sufficient to alter DR5 disulfide bonding alone, other ER stressors may perturb DR5 disulfide bonding, particularly if combined with agents that override the ISR. Consistent with this, treatment of MDA-MB-468 cells with dFtcyDTDO altered DR5 levels and disulfide bonding, while Tunicamycin increased DR5 levels without altering its mobility, and PERKi alone had no discernable effect (Fig. 3B). Tunicamycin + PERKi induced a partial shift in DR5 mobility, similar to that seen with dFtcyDTDO treatment, and this shift was associated with higher Caspase 8 cleavage and more numerous Caspase 3 cleavage products. Unlike DR5, DR4 is *N*-glycosylated. DR4 levels were not changed under any of the conditions, but DR4 mobility was increased by Tunicamycin, presumably due to its deglycosylation.

**Fig. 3:**
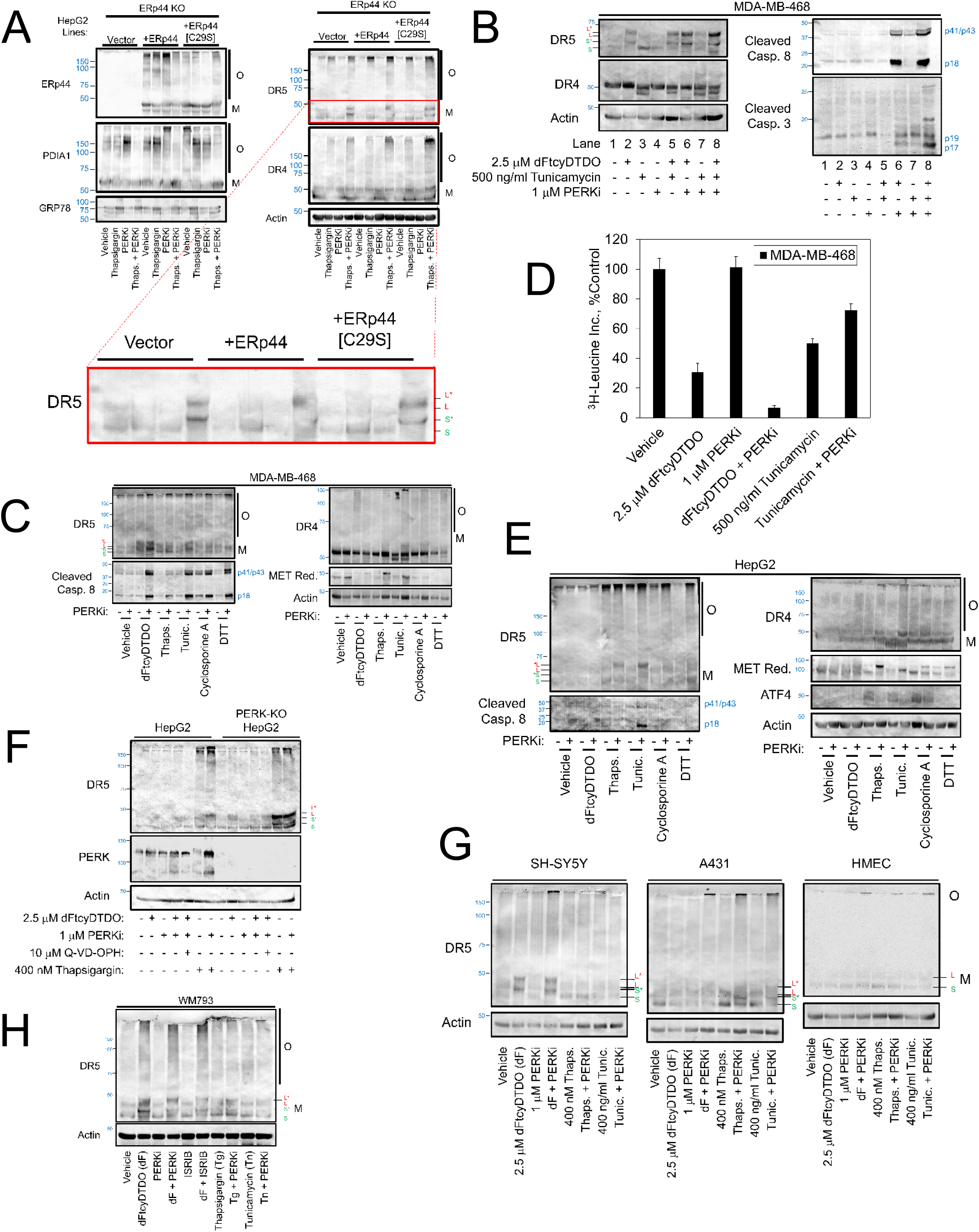
A variety of ER stressors alter DR5 disulfide bonding. A, upper panel. ERp44-deficient HepG2 cells into which vector, wild type or catalytically null ERp44 were reintroduced were treated as indicated for 24 h and analyzed by non-reducing immunoblot. A, lower panel. Expanded region of the DR5 immunoblot showing altered DR5 disulfide bonding in the Thapsigargin/PERKi combination treatment. B. Non-reducing immunoblot of MDA-MB-468 cells treated as indicated for 24 h. C. MDA-MB-468 cells were treated as indicated for 24 h and subjected to non-reducing (DR5, Cleaved Caspase 3, DR4, Actin) or reducing (MET) immunoblot analysis. D. Protein synthesis assays of cells pre-treated for 24 h as indicated before protein synthesis was measured by ^3^H-Leucine incorporation over a 2 h pulse. Data are plotted as the average (N = 6), with error bars representing standard deviation. E. HepG2 cells were treated as indicated for 24 h and subjected to non-reducing (DR5, Cleaved Caspase 3, DR4, Actin) or reducing (MET) immunoblot analysis. F. Control and PERK knockout HepG2 cells were treated for 24 h as indicated and subjected to non-reducing immunoblot analysis. G. Non-reducing immunoblot analysis of the indicated cell lines treated as specified for 24 h. Note the DR5 oligomerization in A431 cells when Thapsigargin, Tunicamycin, or dFtcyDTDO are combined with PERK inhibition. H. Non-reducing immunoblot analysis of WM793 cells treated as specified for 24 h. Unless otherwise specified, the following concentrations of compounds were used in A-H above: dFtcyDTDO (2.5 μM), Thapsigargin (400 nM), Tunicamycin (500 ng/ml), Cyclosporine A (10 μM), Dithiothreitol (DTT; (2.5 mM)), ISRIB (200 nM), or PERKi (1μM). O and M represent Oligomeric and Monomeric protein isoforms in panels A, C, E, G, and H.

Analysis of the effects of other ER stressors on MDA-MB-468 cells indicated that while PERKi alone had little effect on Caspase 8 cleavage, PERKi potentiated induction of Caspase 8 cleavage by Thapsigargin, Tunicamycin, Cyclosporine A, and Dithiothreitol (Fig. 3C). Increased Caspase 8 cleavage correlated with DR5 oligomerization, and in some cases, reduced mobility of monomeric DR5. Immunoblot of MET under reducing conditions showed that dFtcyDTDO and Thapsigargin induced sERr. Tunicamycin + PERKi induction of sERr is difficult to assess given the potentially offsetting effects of deglycosylation and lack of MET proteolytic processing. PERKi caused greater accumulation of unprocessed MET in combination with Thapsigargin than with dFtcyDTDO. Consistent with this observation, protein synthesis assays showed that PERKi increased protein synthesis in the presence of Tunicamycin, but further decreased protein synthesis in the presence of dFtcyDTDO (Fig. 3D). Analysis of the effects of ER stressors on HepG2 cells showed little effect of PERKi or dFtcyDTDO on DR5 levels, while PERKi caused DR5 mobility shifts when combined with Thapsigargin or Tunicamycin (Fig. 3E). Thapsigargin + PERKi caused sERr as assessed by MET processing. Levels of the PERK downstream effector ATF4 increased in response to Thapsigargin, Tunicamycin, and Cyclosporine A. In each case ATF4 upregulation was partially reversed by PERKi.

We next compared ER stress responses observed in PERK knockout and control HepG2 cells. In control cells Thapsigargin increased monomeric and oligomeric DR5 levels and PERKi-cotreatment caused upshifting of the long DR5 isoform and increased DR5 oligomerization (Fig. 3F). As expected, PERKi had no discernable effect on the levels or electrophoretic mobility of DR5 in the PERK knockout cells. We consistently observed a smaller band recognized by the PERK antibody in the presence of PERKi + ER stressors. This likely results from Caspase cleavage of PERK since it is decreased by Caspase inhibitor Q-VD-OPH. Due to differences in the responses of MDA-MB-468 and HepG2 cells to the various ER stressors, we examined ER stress-induced changes in DR5 electrophoretic mobility under non-reducing conditions in neuroblastoma (SH-SY5Y), cervical carcinoma (A431), and human mammary epithelial (HMEC) cells. In SH-SY5Y cells, dFtcyDTDO induced an upward DR5 shift. dFtcyDTDO + PERKi did not further slow DR5 mobility, but increased DR5 oligomerization (Fig. 3G). DR5 mobility and oligomerization were most strongly affected by PERKi combined with Thapsigargin or Tunicamycin in the A431 cells. HMECs exhibited the smallest effect of any of the ER stressors on monomeric DR5 levels, but exhibited a low level of DR5 oligomerization when ER stressors were combined with PERKi. Similar analyses in WM793 melanoma cells showed that dFtcyDTDO alone induced DR5 shifts under non-reducing conditions that were not further potentiated by PERKi or ISRIB (Fig. 3H). Thapsigargin, but not Tunicamycin, induced partial DR5 shifts that were accentuated by PERKi. Together, the findings in Figs. 1 and 2 reveal that changes in DR5 disulfide bonding caused by ER stressors differ among various cancer and non-transformed cell lines and that in some cases ER stress alone is sufficient to alter monomeric DR5 mobility and induce its oligomerization, while in other cases, these effects are potentiated by PERKi.

### EGFR overexpression elevates DDA-induced accumulation of mis-disulfide bonded, monomeric DR5

Given previous observations that DDAs downregulate HER-family proteins [16], and that EGFR overexpression sensitizes cells to DDA cytotoxic effects [17], we examined if DDAs differentially perturb DR5 disulfide bonding and levels in various cancer lines versus non-transformed cells. The DDAs dMtcyDTDO and dFtcyDTDO did not increase DR5 levels in non-transformed MCF10A mammary epithelial cells, HaCaT human keratinocytes, or the T47D luminal breast cancer cell line (Fig. 4A). In contrast, the DDAs induced robust increases in DR5 expression in the MDA-MB-468 and HCC1937 triple-negative breast cancer cell lines. Similarly, while dFtcyDTDO, dFtcyDTDO + PERKi, and Tunicamycin + PERKi reduced DR5 mobility in MDA-MB-468 cells (Fig. 4B), only Tunicamycin + PERKi increased DR5 levels and decreased its mobility in HMECs. Since MDA-MB-468 cells express high EGFR levels [18], we examined if EGFR overexpression is sufficient to confer sensitivity of DR5 to DDA-induced changes in disulfide bonding in MCF10A cells. dFtcyDTDO decreased DR5 mobility in the EGFR overexpressing cells, but not the vector control cells (Fig. 4C). ER stressor Cyclosporine A did not induce this effect. Analysis of the disulfide bonding status of the DDA targets AGR2, ERp44, and PDIA1 showed that EGFR overexpression increased levels of disulfide-bonded oligomers of these PDIs as observed previously [7]. AGR2 is secreted by some cells [19–22], so we examined if dFtcyDTDO or Cyclosporine A caused AGR2 secretion. As expected [23], Cyclosporine A caused Cyclophilin B secretion, but AGR2 secretion was not observed under these conditions.

**Fig. 4:**
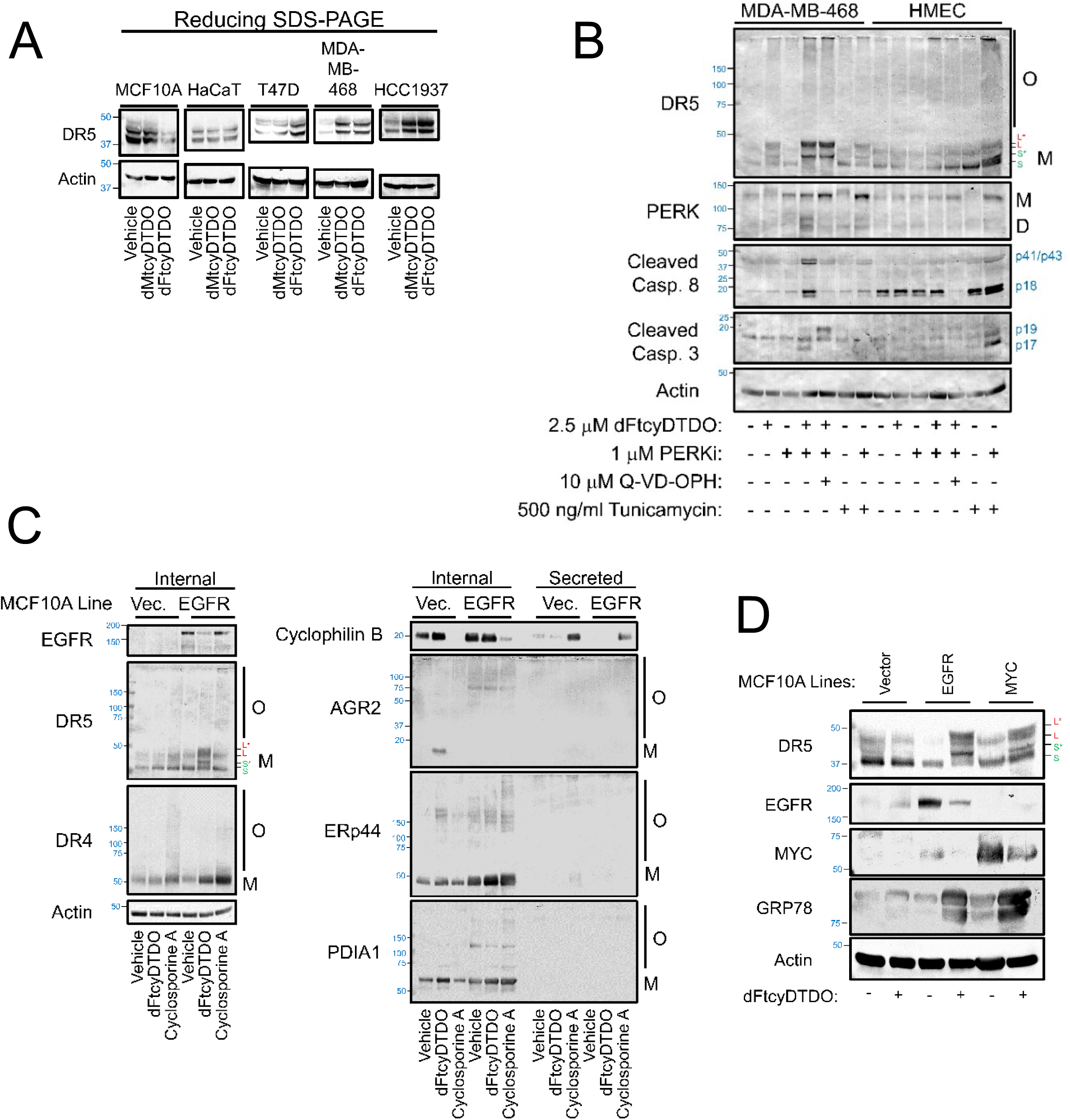
DDA upregulation of DR5 occurs in breast cancer cells or mammary epithelial cells overexpressing MYC or EGFR. A. The indicated cell lines were treated for 24 h with 2.5 μM dMtcyDTDO or dFtcyDTDO and analyzed by immunoblot under reducing conditions. B. MDA-MB-468 cells or Human Mammary Epithelial Cells (HMEC) were treated for 24 h as indicated and subjected to non-reducing immunoblot. D may represent PERK degradation products produced by Caspases. C. MCF10A cells engineered to overexpress EGFR or the corresponding vector control line were treated as indicated for 24 h with 2.5 μM dFtcyDTDO, 10 μM Cyclosporine A, or vehicle. The medium was collected and concentrated for analysis of secreted proteins and the cell extracts were analyzed for internal proteins. Immunoblot analysis was performed under non-reducing conditions. O and M represent Oligomeric and Monomeric protein isoforms in panels B and C. D. MCF10A cells engineered to overexpress EGFR or MYC were treated for 24 h with 2.5 μM dFtcyDTDO or vehicle and cells were analyzed by non-reducing immunoblot. Bands shown represent monomeric protein isoforms.

Since we previously observed that as with EGFR, MYC overexpression sensitizes cells to DDA-driven apoptosis [7], we examined the effects of MYC on DR5 disulfide bonding. dFtcyDTDO shifted DR5 mobility in MYC overexpressing cells, albeit not to the extent observed with EGFR overexpression (Fig. 4D). GRP78 immunoblot indicated that dFtcyDTDO induced a stronger ER stress response in the EGFR and MYC overexpressing cells as compared with the vector control. We previously showed that DDAs selectively upregulate DR5 in a subset of cancer cells and oncogene transformed epithelial lines [7, 8]. The results in Fig. 4 show that this DR5 upregulation by ER stressors is associated with changes in the disulfide bonding of the monomeric forms of DR5, and in some cases is associated with disulfide-mediated DR5 oligomerization.

### DR5 levels and oligomerization states are altered by perturbation of DR5 auto-inhibitory domain disulfides

A recent study showed that the disulfide bond-rich extracellular domain of DR5 serves to prevent receptor oligomerization and pro-apoptotic signaling in the absence of its ligand TRAIL [24]. A more recent article narrowed down the DR5 autoinhibitory domain to a positive patch involving residues R154, K155, and R157 [25]. Based on a crystal structure [26], these basic residues share a common orientation due to two disulfide bonds, C156-C170 and C139-153 (Fig. 5A). We hypothesized that loss of these two disulfide bonds may disrupt the auto-inhibitory domain, culminating in DR5 clustering and activation of Caspase 8-Caspase 3-mediated apoptosis independently of TRAIL or DDA treatment. This hypothesis was tested by doxycycline-inducible expression of wild type and disulfide-defective DR5 point mutants. High-level inducible expression of the long form of wild type DR5 required induction by doxycycline combined with DDA treatment as described previously [8], while mutation of one or both of the Cys residues of the C160-C178 disulfide bond conferred high level DR5 expression in the absence of DDA treatment, as did the C81S mutation (Fig. 5B). Interestingly, DDA treatment still caused an upward shift in these mutants under reducing conditions suggesting that DDAs may disrupt multiple DR5 disulfide bonds. The DR5 disulfide-defective mutants, but not wild type DR5, also caused formation of high molecular weight DR4 oligomers in the absence of DDA treatment suggesting that endogenous DR4 may co-aggregate with ectopic, mis-disulfide bonded DR5 oligomers.

**Fig. 5:**
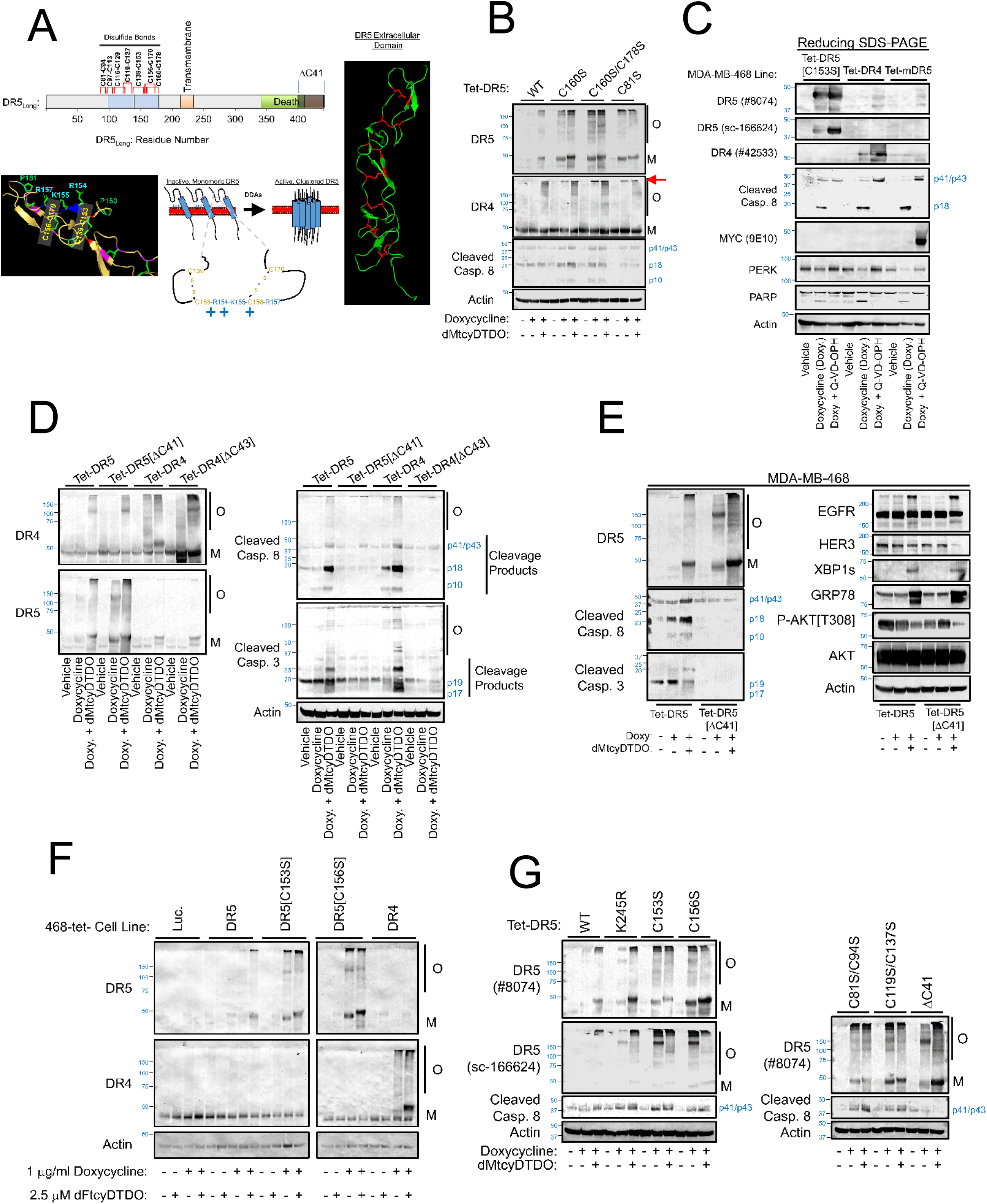
Genetic disruption of multiple DR5 disulfide bonds induces its stabilization and pro-apoptotic signaling. A. Structural model of DR5 showing its disulfide bonds, and the positive patch autoinhibitory domain described in the literature. B. Non-reducing immunoblot analysis of MDA-MB-468 cells engineered with doxycycline-inducible expression of wild type (WT) DR5 or the indicated Cys to Ser disulfide bond mutants. Cells were treated as indicated for 24 h with 1 μg/ml doxycycline and 2.5 μM dMtcyDTDO. The red arrow denotes DR4 oligomers that coincide with DR5 oligomerization. C. Reducing immunoblot analysis of the indicated MDA-MB-468 stable cell lines. Cells were treated for 24 h as specified with 1 μg/ml doxycycline or doxycycline + 10 μM Q-VD-OPH. The catalog numbers of DR5 and DR4 antibodies are shown. D. Non-reducing immunoblot analysis of the indicated MDA-MB-468 cell lines with doxycycline-inducible expression of wild type DR4 and DR5, and DR4 and DR5 C-terminal deletion constructs defective in apoptotic signaling. Cells were treated for 24 h as specified with 1 μg/ml doxycycline or doxycycline + 2.5 μM dMtcyDTDO. E. Non-reducing immunoblot analysis of the indicated MDA-MB-468 cell lines with doxycycline-inducible expression of wild type and apoptosis-defective DR5. Cells were treated for 24 h as specified with 1 μg/ml doxycycline or doxycycline + 2.5 μM dMtcyDTDO. F. Non-reducing immunoblot analysis of the indicated MDA-MB-468 doxycycline-inducible stable cell lines. Cells were treated for 24 h as indicated. G. Non-reducing immunoblot analysis of the indicated MDA-MB-468 cell lines with doxycycline-inducible expression of wild type and apoptosis-defective DR5. Cells were treated for 24 h as specified with 1 μg/ml doxycycline or doxycycline + 2.5 μM dMtcyDTDO. O and M represent Oligomeric and Monomeric protein isoforms in panels B and D-G.

We next examined if Caspase activation limits expression of DR5[C153S], DR4, or the murine TRAIL receptor (mDR5) by inhibiting Caspases with Q-VD-OPH [27]. Q-VD-OPH increased the inducible expression of all three receptors and prevented formation of the p18 fragment of Caspase 8, but not the p41/p43 fragment (Fig. 5C). This likely indicates that Q-VD-OPH does not inhibit receptor-driven Caspase 8 autocleavage, but inhibits the previously described [28] Caspase 3 cleavage of Caspase 8. We further examined the relationship between DR5 and DR4 oligomerization and Caspase activation using C-terminal DR5 or DR4 deletion constructs since such mutants were previously shown incapable of coupling to Caspase 8 activation [29]. Doxycycline and doxycycline + dMtcyDTDO produced similar effects on the levels of the wild type and mutant DR5 and DR4, although the mutants were incapable of activating the Caspase 8-Caspase 3 cascade (Fig. 5D). Since DDAs cause ER stress, we examined if ER stress is independent of DR5-mediated Caspase activation. dMtcyDTDO upregulated ER stress markers, decreased AKT phosphorylation, and increased disulfide-mediated EGFR oligomerization irrespective of Caspase activation (Fig. 5E).

DR5 mutants lacking the disulfide bonds that form the positive patch exhibit high expression and oligomerization in the absence of DDA treatment, unlike wild type DR5 (Fig. 5F). Upregulation of DR5 by disruption of positive patch Cys residues was observed with two antibodies that recognize the C-terminal (#8074) or N-terminal (sc-166624) regions of DR5, although the latter antibody exhibited a strong binding preference for oligomeric DR5 isoforms over monomeric DR5 (Fig. 5G). The PhosphoSite database ((Phosphosite.org) lists K245 as a major site of DR5 ubiquitination. Mutation of this site to Arg modestly increased receptor levels in the doxycycline and doxycycline + DDA treated samples, but did not mimic the ability of the C-S mutations to exhibit high level expression in the absence of DDA treatment. In summary, the results in Fig. 5 indicate that individual mutation of several different DR5 disulfides, including the positive patch disulfides, is sufficient for high level expression of DR5 and activation of Caspases independent of DDA treatment or ER stress. Further, stabilization of mis-disulfide bonded DR5 does not require Caspase activation, but, activation of Caspase 8 by mis-disulfide bonded DR5 requires its C-terminus that is necessary for DISC formation [29].

### DDAs activate autophagy and autophagy inhibitors potentiate DDA-induced DR5 accumulation

Proteasomal degradation (ERAD) and autophagy are important modes for the disposal of misfolded proteins. However, the fate of mis-disulfide bonded, aggregated proteins in the secretory pathway is underexplored. Since DDAs induce ER stress [17], and ER stress frequently activates autophagy [30], we examined if DDAs stimulate autophagy. dFtcyDTDO treatment induced an upward DR5 shift in a concentration-dependent manner. GPR78 expression and autophagy marker LC3 lipidation were both increased to maximal levels at the lowest dFtcyDTDO concentration tested (Fig. 6A). Treatment with the autophagy/lysosome inhibitor Bafilomycin A1 increased levels of monomeric and oligomeric DR5 isoforms in the absence of DDA treatment (Fig. 6B). Similar studies employing the autophagy/lysosome inhibitor Chloroquine showed increased DR5 levels and accumulation of oligomeric EGFR compared with vehicle treatment. Combining dFtcyDTDO with Chloroquine or PERKi increased DR5 levels and Caspase 8 cleavage over that observed with dFtcyDTDO alone (Fig. 6C). Cell viability assays of cells treated as in Fig. 6C showed that combining dFtcyDTDO with either PERKi or Chloroquine reduced viability more than dFtcyDTDO, and combining the three agents decreased viability the most (Fig. 6D). An inhibitor of the autophagy PI3-kinase VPS34 (VPS34i, [31]) increased DR5 expression to a similar extent as Bafilomycin, and combining dFtcyDTDO with either VPS34i or Bafilomycin increased expression of monomeric DR5 more than each individual treatment (Fig. 6E). Of these treatments, only dFtcyDTDO or the dFtcyDTDO-containing treatments upshifted monomeric DR5. The strong accumulation of upshifted monomeric DR5 caused by autophagy inhibitors suggests that autophagy plays a role in degrading mis-disulfide bonded DR5.

**Fig. 6:**
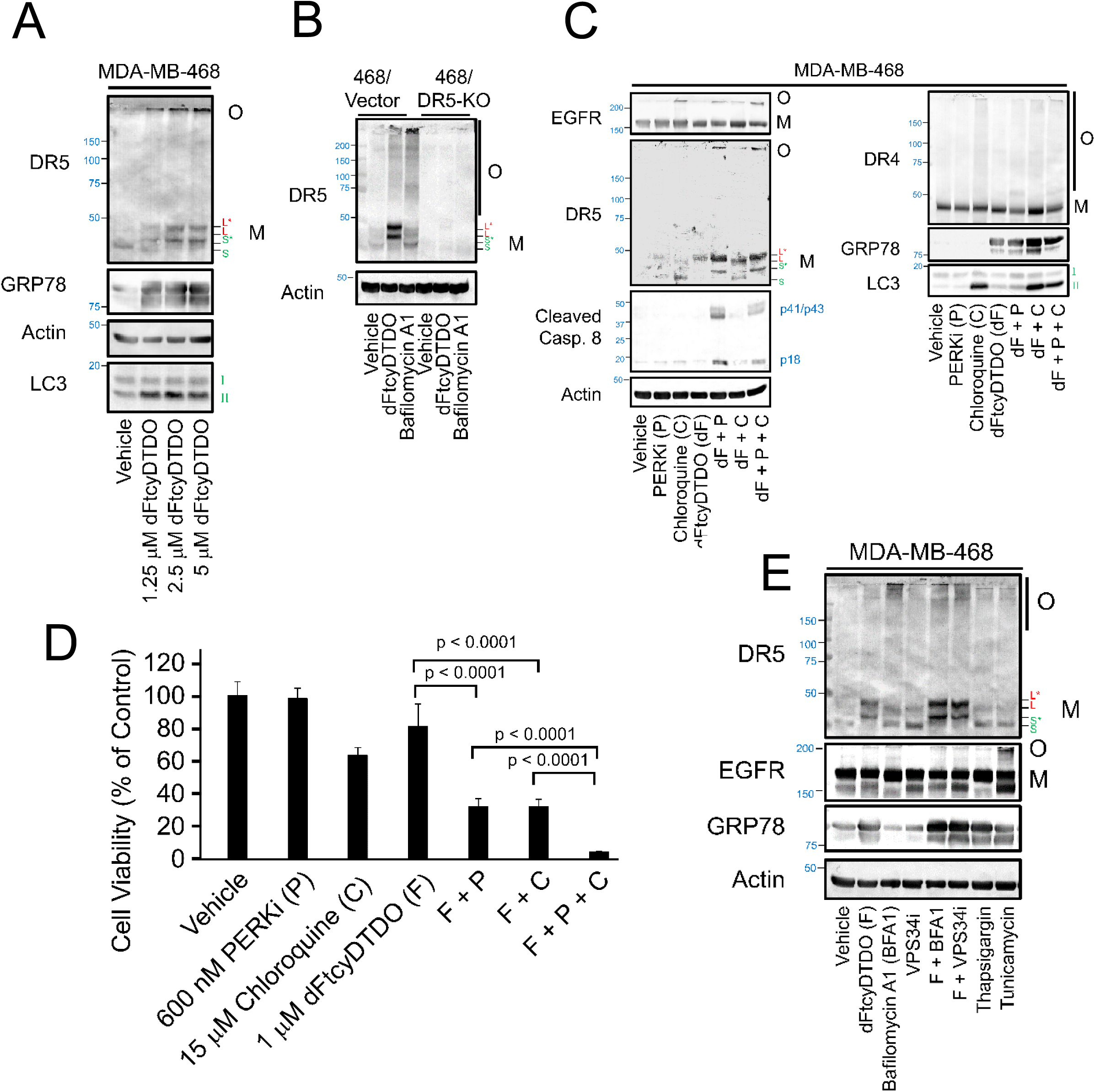
DDAs activate autophagy and inhibitors of autophagy/lysosomal degradation upregulate DR5. A. Non-reducing immunoblot analysis of MDA-MB-468 cells treated as indicated for 24 h. B. Non-reducing immunoblot analysis of vector control or DR5 knockout MDA-MB-468 cells treated with 2.5 μM dFtcyDTDO or 1 μM Bafilomycin A1 for 24 h. C. Non-reducing immunoblot analysis of MDA-MB-468 cells treated for 24 h as indicated with 1 μM PERKi (P), 15 μM Chloroquine (C), or 2.5 μM dFtcyDTDO (dF). D. MTT Cell viability assay of MDA-MB-468 cells treated as indicated for 72 h. Data are plotted as the average (N = 6), with error bars representing standard deviation. E. Non-reducing immunoblot analysis of MDA-MB-468 cells treated for 24 h as indicated with 2.5 μM dFtcyDTDO (F) or 1 μM Bafilomycin A1 (BFA1), VPS34 inhibitor (VPS34i), 400 nM Thapsigargin, or 500 ng/ml Tunicamycin. O and M represent Oligomeric and Monomeric protein isoforms in panels A, B, C, and E.

### DDAs upregulate DR5 paralog DcR2

The pro-apoptotic TRAIL receptors DR4 and DR5 share conserved disulfide-rich domains with the TRAIL decoy receptors DcR1 and DcR2 that bind TRAIL, but cannot activate Caspases. Specifically, the DR5 autoinhibitory domain is largely conserved with DR4, DcR1, and DcR2 (Fig. 7A). We considered that DDAs might stabilize DcR1 or DcR2 in a similar manner as DR5. Since we previously found that the prolyl isomerase inhibitor Cyclosporine A potentiated DDA cytotoxic effects [9], we examined Cyclosporine A effects on the levels of decoy receptors. We did not detect DcR1 in the cell lines examined, but observed upregulation of DcR2 after dFtcyDTDO treatment (Fig. 7B). We also observed that Cyclosporine A, but not FK606, which inhibits a different family of prolyl isomerases, decreased DDA upregulation of DcR2, but not DR5. dFtcyDTDO increased DcR2 levels more at low concentrations than high concentrations in some experiments, but irrespective of the pattern of DcR2 upregulation by dFtcyDTDO, it was blocked by co-treatment with Cyclosporine A (Fig. 7B-D). Quantitation of band intensities revealed that Cyclosporine A treatment potentiated the effects of low (310 nM) dFtcyDTDO concentration on DR5 oligomerization (Fig. 7E, left panel) and decreased DcR2 upregulation by dFtcyDTDO (Fig. 7E, right panel).

**Fig. 7:**
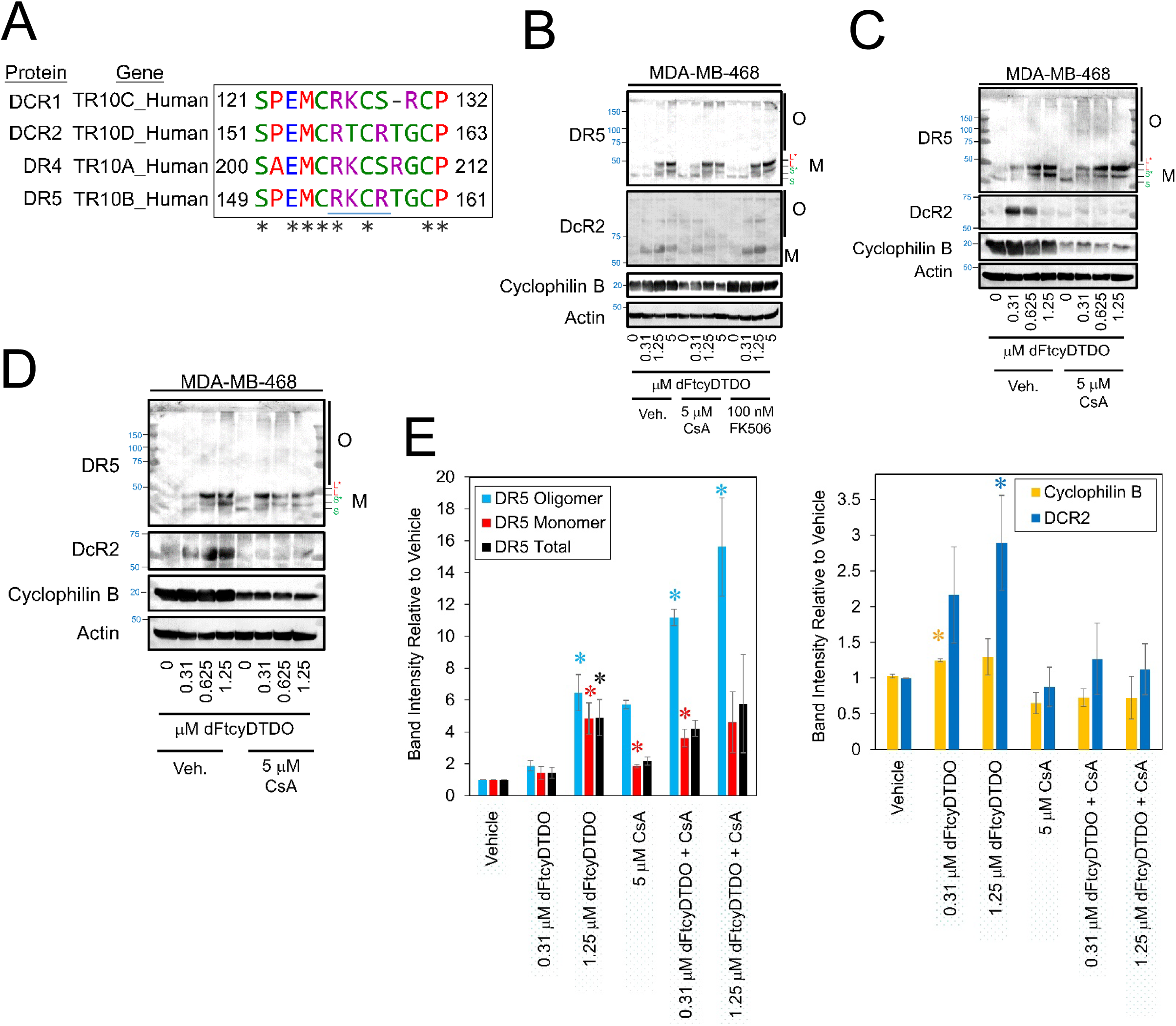
DDAs upregulate TRAIL Decoy Receptor 2, an effect overridden by Cyclosporine A. A. Sequence alignment of the putative autoinhibitory motifs of DR4, DR5, DCR1, and DCR2. B. Non-reducing immunoblot analysis of MDA-MB-468 cells treated for 24 h with the indicated concentrations of dFtcyDTDO, combined with DMSO vehicle, 5 μM Cyclosporine A or 100 nM FK506. C. Non-reducing immunoblot analysis of MDA-MB-468 cells treated for 24 h with the indicated concentrations of dFtcyDTDO, combined with DMSO vehicle or 5 μM Cyclosporine A. D. Non-reducing immunoblot analysis of MDA-MB-468 cells treated for 24 h with the indicated concentrations of dFtcyDTDO, combined with DMSO vehicle or 5 μM Cyclosporine A. Data are plotted as the average (N = 3), with error bars representing standard error. Asterisks denote *p* < 0.05 compared to control using Student’s unpaired *t*-test. E. Densitometry analysis of the relative levels of total, monomeric, and oligomeric forms of DR5 (left panel) or total levels of Cyclophilin B or DCR2 (right panel) from panels 7B-D. O and M represent Oligomeric and Monomeric protein isoforms in panels B-D.

### DR5 bonding status influences antibody recognition, but not trafficking to the cell surface

Since sERr prevents some transmembrane receptors from reaching the cell surface [12] and DR5 was shown to be activated in the Golgi by binding to aggregated proteins [32], we examined DR4 and DR5 cell surface labeling by flow cytometry. Using the Clone DJR2-4 (7-8) antibody for flow cytometry analyses, Doxycycline induction of the wild type, C153S, and C156S mutants showed DR5 increased cell surface labeling, however co-treatment with dFtcyDTDO decreased apparent DR5 surface localization (Fig. 8A, upper panel). DR4 flow cytometry studies showed that doxycycline increased DR4 surface levels and this was not altered by dFtcyDTDO co-treatment. Since recognition of DR5 by the flow cytometry antibody could be hindered by changes in DR5 disulfide bonding, we examined levels of DR5 and its disulfide bonding mutants using three different commercially available antibodies (Fig. 8B). The #8074 antibody directed against the cytoplasmic, C-terminal portion of DR5 recognized all of the DR5 proteins, including the monomeric and oligomeric forms of DR5. As shown here (Fig. 5) and elsewhere [8], wild type DR5 was only maximally expressed in cells induced with doxycycline and treated with DDAs. The sc166624 DR5 antibody directed against the N-terminal cysteine-rich region preferentially recognized the oligomeric forms of DR. In contrast, Clone DJR2-4 (7-8) directed toward the N-terminal cysteine-rich portion of DR5 did not recognize the C119S/C137S DR5 mutant, but bound the C153S, C156S, C160S, and C160S/C178S DR5 mutants. However, dFtcyDTDO treatment ablated DR5 recognition by this antibody. These observations suggest that binding of Clone DJR2-4 (7-8) antibody is sensitive to DR5 disulfide bonding, which is altered by DDA treatment. Since this antibody is commonly used for DR5 labeling in flow cytometry studies and multiple ER stressors, including DDAs, alter DR5 disulfide bonding, lack of signal with this antibody may be indicative of changes in DR5 disulfide bonding rather than DR5 downregulation or internalization.

**Fig. 8:**
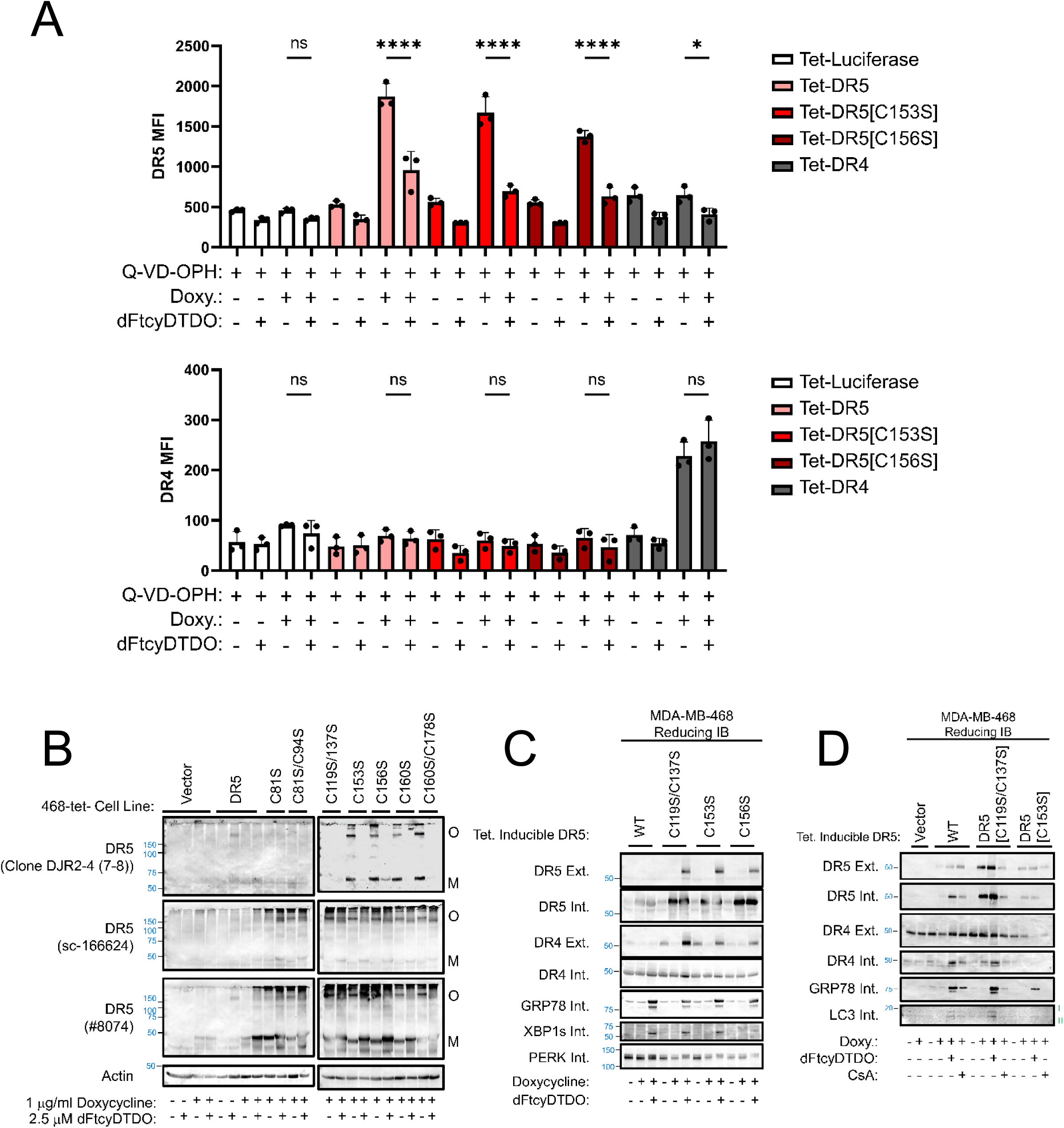
Effects of altered disulfide bonding on DR5 cell surface localization and antibody recognition. A. Flow cytometry analysis of the indicated doxycycline-inducible MDA-MB-468 stable cell lines with an antibody to DR5 (Clone DJR2-4 (7-8)) (top panel) or DR4 (bottom panel). Prior to analysis, cells were treated for 24 h as indicated with 10 μM Q-VD-OPH, 1 μg/ml doxycycline, or 2.5 μM dFtcyDTDO. Dots represent the average values from three independent biological replicates performed in triplicate. * Represents p < 0.05, **** represents p < 0.0001, and ns represents not significant (p > 0.05). B. Non-reducing immunoblot analysis of the indicated MDA-MB-468 stable cell lines treated for 24 h as specified. Note the alternate staining patterns observed with different DR5 antibodies. O and M represent Oligomeric and Monomeric protein isoforms. C. The indicated MDA-MB-468 stable cell lines were treated for 24 h as indicated with 1 μg/ml doxycycline and 2.5 μM dFtcyDTDO and subjected to cell surface protein biotin labeling. Cell surface proteins (External; Ext.) were affinity purified with Streptavidin-agarose, and the unlabeled flow-through (Internal; Int.) proteins were also collected. Both fractions were analyzed by non-reducing immunoblot using the indicated antibodies. D. Cell surface protein labeling experiment as in panel C except that cell lines were treated with the indicated combinations of 1 μg/ml doxycycline, 2.5 μM dFtcyDTDO, and 10 μM Cyclosporine A.

Previous work showed that DDA treatment increased surface localization of DR5 as measured by biotin labeling [7]. The same approach was used to examine the localization of the DR5 mutants to the cell surface. The results showed that the C119S/C137S, C153S, and C156S mutants trafficked to the cell surface, particularly in the context of DDA treatment (Fig. 8C). Under these conditions, expression of DR5 disulfide bonding mutants did not elicit an ER stress response as indicated by the markers GRP78, XBP1s, and PERK activation (phosphorylation). However, dFtcyDTDO activated all these indicators of ER stress. A similar cell surface biotinylation experiment showed trafficking of wild type and mutant DR5 to the cell surface. Surface localization of the C119S/C137S and C153S DR5 mutants occurred in both the presence or absence of dFtcyDTDO or the ER stress inducer Cyclosporine A (Fig. 8D). Expression of mis-disulfide bonded DR5 mutants did not upregulate the ER stress or autophagy markers GRP78, or LC3, respectively, but DDA treatment upregulated both markers. Together, these results indicate that under conditions where DR5 disulfide bonding is perturbed by either mutagenesis or DDA treatment, DR5 traffics to the cell surface. This is consistent with previous work showing that DDA treatment increases cancer cell sensitivity to the DR4/5 ligand TRAIL [8].

### DDA safety and identification of a metabolically stable DDA analog

Previous work with the DDAs RBF3 [16] and tcyDTDO [8] did not reveal evidence of toxicity under conditions in which they induce the death of primary and metastatic breast cancer cells in mouse models. We also examined the metabolism of the DDA tcyDTDO by liver microsomes [9], but did not present analyses of the stability of dMtcyDTDO and dFtcyDTDO toward metabolism in liver and intestinal microsomes. Recent work demonstrated the activity of both dMtcyDTDO and dFtcyDTDO in mouse models of breast cancer [33], but did not examine the effects of these compounds on normal tissues such as the liver or hematopoietic cells. Examination of breast tumor tissue from mice treated with vehicle or 10 mg/kg dMtcyDTDO showed extensive death of tumor tissue in the dMtcyDTDO-treated, but not the vehicle-treated mice (Fig. 9A, upper panels). Liver tissues from vehicle or dMtcyDTDO-treated mice were indistinguishable (Fig. 9A, lower panels). Analysis of complete blood cell counts from tumor-bearing mice treated with vehicle or 10 mg/ml dMtcyDTDO were in the normal range for healthy mice (Supplemental Table: S1). The apparent decrease in platelets across the samples was likely due to partial clotting prior to analysis.

**Fig. 9:**
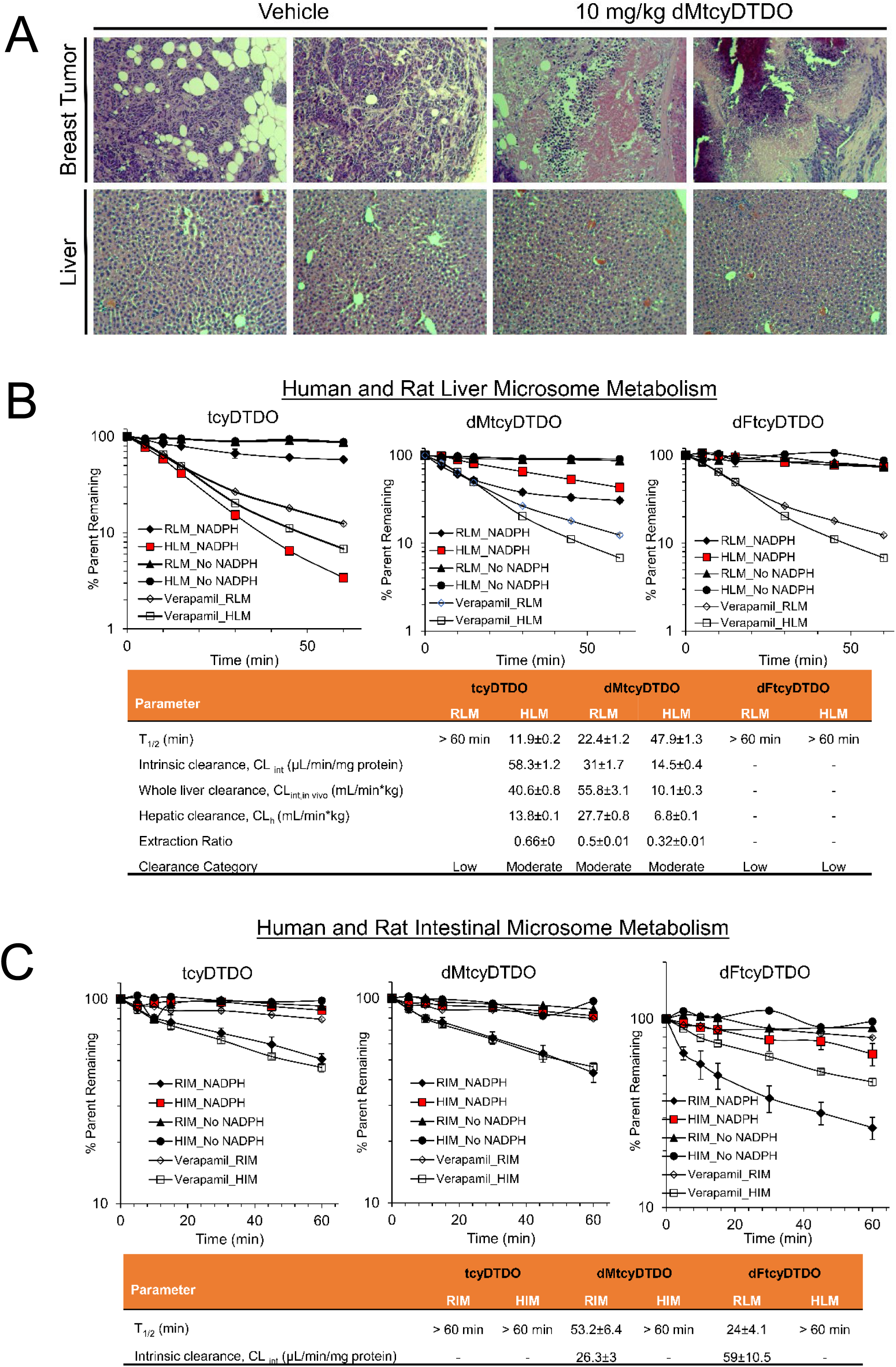
Metabolic stability of select DDAs and lack of DDA effects on liver morphology. A. Hematoxylin and Eosin stained breast tumor (upper panels) and liver tissue samples (lower panels) from mice bearing 012/LVM2/LR10 tumors after treatment with vehicle (peanut oil) or 10 mg/kg dMtcyDTDO by oral gavage for 20 days. B. Stability of tcyDTDO, dMtcyDTDO, or dFtcyDTDO metabolism in rat or human liver microsomes in the presence or absence of 1 mM NADPH. Verapamil serves as a positive control. C. Stability of tcyDTDO, dMtcyDTDO metabolism, or dFtcyDTDO in rat or human intestinal microsomes in the presence or absence of 1 mM NADPH. Verapamil serves as a positive control. Data points are plotted as the average (N = 3), with error bars representing standard deviation.

Stability studies in human liver microsomes supplemented with NADPH showed that FtcyDTDO was metabolically stable (t_1/2_ > 60 min), while tcyDTDO, and dMtcyDTDO metabolized by phase I enzymes with half-lives of 11.9, and 47.9 min, respectively. Similar studies employing human intestinal microsomes showed that the half-life of all three DDA compounds exceeded 60 min. Although more thorough analysis of these DDAs is needed, studies performed to date indicate that dFtcyDTDO has a more favorable metabolic stability profile than tcyDTDO or dMtcyDTDO.

## Discussion

Previous work showed that the DDA compounds induce ER stress, which is associated with high-level DR5 expression and disulfide-mediated oligomerization [8, 17, 34]. Further, mutational disruption of a subset of DR5 disulfide bonds was demonstrated to stabilize DR5 and trigger disulfide-mediated DR5 oligomerization. However, the relationship between the DDA-induced ER stress response and DDA effects on DR5 were unexplored. The results presented here show that individual mutational disruption of a total of five of the seven DR5 disulfide bonds each causes the same DR5 stabilization, including the disulfides within the previously described auto-inhibitory domain [24, 25]. The observation that DDAs still induce a mobility shift of monomeric disulfide point mutants of DR5 suggests that while loss of individual disulfide bonds is sufficient to stabilize DR5 and promote pro-apoptotic signaling, DDAs disrupt multiple DR5 disulfide bonds.

Numerous studies have demonstrated transcriptional regulation of DR5 through a PERK-ATF4-CHOP-DR5 pathway [35, 36]. In this model, PERK inhibition is predicted to block DR5 upregulation by ER stress. Our previous studies in breast cancer lines found little effect of knocking out or overexpressing CHOP on DR5 levels, suggesting that other DR5 regulatory mechanisms exist [8]. We found here that PERK inhibition potentiated DR5 pro-apoptotic signaling, but only modestly increased DR5 levels. A report showed that DR5 is activated by binding to misfolded proteins in the Golgi [32], so PERKi might elevate DR5 oligomerization through this mechanism. Alternatively, overriding ISR may activate DR5 by increasing protein folding flux under ER stress conditions and surpass the ability of the DDA targets ERp44, PDIA1, and AGR2 to catalyze disulfide bond formation or their functions in protein folding checkpoints. A recent report showed that breast cancer metastases exhibit elevated ER stress and are responsive to a new, more selective PERK inhibitor [37]. This along with our previous study showing DDA activity against breast cancer metastases [8] provides a rationale for DDA/PERKi combination therapy for the treatment of metastatic breast cancer. Likewise, the observation that Cyclosporine A blocks DDA-induced upregulation of DCR2 further supports the previous contention [9] that DDA/Cyclosporine co-treatment may exhibit enhanced efficacy against breast malignancies.

Overexpression of the EGFR or MYC oncoproteins was shown to sensitize cells to DDA cytotoxic effects, but the underlying mechanisms were not investigated [8]. Results presented here show that EGFR or MYC overexpression permits DDA perturbation of DR5 disulfide bonding that is not observed in vector control non-transformed MCF10A mammary epithelial cells. This may partially explain the ability of DDAs to mimic the cancer-specific cytotoxic effects of the DR5/4 ligand TRAIL. Interestingly, expression of disulfide bonding mutants of DR5 does not trigger an ER stress response or activate autophagy. Further, mis-disulfide bonded DR5 traffics to the cell surface, consistent with the previous observation that DDAs synergize with TRAIL to kill cancer cells [8]. The present results extend previous work on the relationship between ER stress and DR5 activation by showing that in addition to DDAs, other ER stressors, including Tunicamycin and Thapsigargin, can alter DR5 disulfide bonding in a manner that is potentiated in some cases by PERKi co-treatment. A subject of ongoing investigation is why ER stressors that alter DR5 disulfide bonding do not have the same effect on its paralog DR4. This may relate to the different N- and O-glycosylation patterns observed for these receptors [38–41], or regulation of DR4 and DR5 stabilities by different E3 ubiquitin ligases [42–44]. Together, our work [7, 8, 17] and that of others [32], suggest that DR5 has evolved as a direct sensor and effector of ER stress/protein misfolding and that ER stress can activate DR5 through transcriptional mechanisms and at the protein level through altered DR5 disulfide bonding and DR5 binding to misfolded proteins. Importantly, DR5 exhibits a TRAIL-independent gain of function under these conditions that inactivate a wide variety of other transmembrane oncoproteins, including EGFR and MET.

DDA studies have been performed largely in breast cancer cell lines, therefore it will be critical to determine the importance of these DDA-driven effects in non-transformed cells and across other tumor types. This DDA cancer selectivity is further supported by the Broad Institute’s Dependency Map (DepMap; depmap.org/portal/). Of the 22 human disulfide isomerases, only four, ERp44, AGR2, AGR3, and TMX1, are considered as “strong dependencies” in the DepMap, and our studies have shown DDAs to inhibit three of these, ERp44, AGR2, and AGR3. It is possible that the client proteins of ERp44, AGR2, and AGR3 vary with tumor type. As an example, out of 17,347 genes in the DepMap CRISPR screen, the colon cancer lines C80, COLO205, LS513, and SNUC4 rank AGR2 as the first, fourth, third, and fifth most important gene, respectively. Interestingly, ERN2 encodes the IRE1α homolog IRE1β whose expression is restricted to Goblet cells. The DepMap lists IRE1β as the top predictor of AGR2 dependence. Two recent reports show that AGR2 functions as an inhibitor of IRE1β that overcomes the cytotoxic effects of this enzyme [45], and that the active site Cys81 of AGR2 is required for IRE1β inhibition [46, 47]. It will be important to determine if DDAs exhibit anticancer activity against tumor lines in which AGR2 is required to prevent IRE1β-mediated cancer cell death. This is an area of active investigation by our team.

More generally, DDAs may serve as important tools for investigating disulfide bonding quality control in the secretory pathway. Significant interest has recently focused on DDAs and similar compounds for their ability to function in transport across cell membranes in thiol-mediated uptake [48] and as redox-sensitive probes [49] indicating the value of six membered cyclic dichalcogenides for diverse biological applications. As an inhibitor of ERp44, DDAs may override two key protein disulfide bonding checkpoints, the retrograde Golgi-ER recycling of secretory proteins discovered by Sitia and Colleagues [50–55], and the selective ER retention mechanism of some receptor tyrosine kinases discovered by Tirosh and Colleagues [12]. Based on these new mechanistic insights and molecular and pharmacological tools, the stage is set to investigate the molecular and biological functions of the non-canonical PDIs, and strongly selective cancer dependencies, ERp44, AGR2, and AGR3.

## Materials and Methods

### Cell culture, preparation of cell extracts, and immunoblot analysis

MCF10A cells were cultured in a humidified incubator set at 37°C containing 5% CO_2_ as described previously [56]. All other cell lines were grown in Dulbecco’s Modified Eagle’s Medium (GE Healthcare Life Sciences, Logan, UT) with 10% fetal bovine serum (10% FBS–DMEM). A431, AsPC1, HaCaT, HCC1937, HepG2, HMEC, MCF10A, MDA-MB-468, SH-SY5Y, and T47D were purchased from American Type Culture Collection (ATCC) (Manassas, VA). SUM149pt was purchased from Applied Biological Materials, Inc. (Richmond, BC, Canada). ERp44 and PERK knockout HepG2 cell lines were described previously [12]. WM793 cells were kindly provided by Dr. W. Douglas Cress, Moffitt Cancer Center. Generation of the MCF10A/Vector, MCF10A/EGFR and MCF10A/MYC cell lines is described in previous work [8]. Derivation of the HCI-012/LVM2/LR10 cell line is described in previous work [8, 9]. Generation of DR5 knock out MDA-MB-468 cells and MDA-MB-468 cells stably expressing DR5 using the Tet-ON system is previously described [8]. Generation of the MDA-MB-468 cells stably expressing CDCP1 or vector control is described in previous work [15].

Cell lysates were prepared as described in a previous publication [57]. Immunoblot analysis was performed employing the following antibodies purchased from Cell Signaling Technology (Beverly, MA) [Akt, #4691; P-Akt[T308], #13038; ATF4, #11815; CDCP1, #13794; Cleaved Caspase 3, #9664; Cleaved Caspase 8, #9496; Cyclophilin B, #43603; DCR2, #8049; DR4, #42533; DR5, #8074; eE2F, #2332; P-eE2F[T56], #2331; GRP78, #3177; HER3, #4754; LC3, #3868; MET, #3127; PARP, #9532; PCSK9, #55728; PDIA1, #3501; PERK, #5683; and XBP1s, #12782], from Santa Cruz Biotechnology (Santa Cruz, CA) [Actin, sc-47778; AGR2/3, sc-376653; AGR3, sc-390940; c-Myc (9E10), sc-40; EGFR, sc-373746; ERp44, sc-393687; PDIA6, sc-365260; and pY99, sc-7020], and from Rockland Immunochemicals, Inc. (Limerick, PA), Streptavidin-Alkaline Phosphatase conjugated (SA-AP), S000-05.

### Quantitative Analysis of Immunoblot Results

Protein levels in immunoblots were quantified using Adobe Photoshop (Berkeley, CA) and ImageJ (NIH, Bethesda, MD), as previously described [58], followed by normalization to Actin as a loading control.

### Materials

Reagents were purchased from the following companies: Tunicamycin and Chloroquine: Sigma-Aldrich (St. Louis, MO); Thapsigargin: AdipoGen (San Diego, CA); Lapatinib: Selleck Chemicals (Houston, TX); Doxycycline: Enzo Life Science (Farmingdale, NY); TORIN1, and dithiothreitol (DTT): TOCRIS Bioscience (Minneapolis, MN); Cyclosporine A (CsA): Biorbyt (Duran, NC); Rapamycin and PERK Inhibitor I (GSK2656157): Calbiochem (Burlington, Massachusetts); Bafilomycin A1, Gefitinib, ISRIB, MK2206, and Q-VD-OPH: Cayman Chemical (Ann Arbor, MI); *N*-ethylmaleimide (NEM): Thermo Fisher Scientific (Grand Island, NY); eEF2K inhibitor (A-484954) and VPS34 inhibitor Vps34-IN-1: MedChemExpress (Monmouth Junction, NY); FK506: InvivoGen (San Diego, CA); b-AP15: MedKoo Biosciences (Chapel Hill, NC).

### Tumor studies and histochemical analysis

012/LVM2/LR10 xenograft tumor studies were carried out in adult female NOD-SCID-γ (NSG) mice, as described in a previous publication [9]. After the development of palpable tumors (approximately 4 mm^3^), mice were randomly assigned to two treatment groups: DMSO (Vehicle) and 10 mg/kg dMtcyDTDO). Mice were treated every weekday for twenty days by intraperitoneal injection, administering 50 µL per injection. At the end of the twenty-day period, tissue samples were collected and fixed in 4% paraformaldehyde/Phosphate-Buffered Saline (PBS), followed by paraffin-embedding, sectioning and staining with hematoxylin and eosin (H&E) by the University of Florida Molecular Pathology Core (https://molecular.pathology.ufl.edu/). Prior to endpoint, peripheral blood was collected by facial vein puncture into EDTA-treated tubes; complete blood cell counts (CBCs) were obtained using an Element HT5 fully automated hematology analyzer (Heska, Loveland, CO).

### Disulfide Bond-mediated Oligomerization

Disulfide bond-mediated oligomerization under non-reducing conditions was analyzed as described in previous work [7].

### Vector Construction

In order to construct the Tet-DR4 expression vector, DR4 (Addgene plasmid #61382) was amplified using the following primers: 5′-TTTTATCGATCACCATGGCGCCCGTCGCCGTCTGG-3′ and 5′-TTTTGGATCCTCACTCCAAGGACACGGCAG-3′ and cloned into the pRetroX-TetOne-Puro vector with a modified cloning site that incorporates Not I, Bcl I, and Cla I sites 5′ to the BamH I site (Clontech, Mountain View, CA, USA).

The initial mutations of C81S, C119S and C160S in DR5 were performed in pcDNA3 with QuikChange mutagenesis and the following primers, respectively: 5′- CCAGCCCCTCAGAGGGATTGAGTCCACCTGGACACCATATC-3- and 5′- GATATGGTGTCCAGGTGGACTCAATCCCTCTGAGGGGCTGG-3′, 5′- GCTTGCGCTGCACCAGGAGTGATTCAGGTGAAGTGG-3′ and 5′- CCACTTCACCTGAATCACTCCTGGTGCAGCGCAAGC-3′ and 5′- CGGAAGTGCCGCACAGGGAGTCCCAGAGGGATGGTCAAGG -3′ and 5′- CCTTGACCATCCCTCTGGGACTCCCTGTGCGGCACTTCCG-3′. Mutations were verified by sequencing. The following primers were used to add a 5′-EcoRI and a 3′-BamHI site to C81S, C119S and C160S DR5 by PCR amplification: 5′- TTTTGAATTCCACCATGGAACAACGGGGACAGAAC-3′ and 5′- TTTTGGATCCTTAATGATGATGATGATGATGGGACATGGCAGAGTCTGC-3′. The C119S and C160S DR5 mutants were subsequently cloned into the EcoRI and BamHI sites of pRetroX-TetOne- Puro vector (Clontech, Mountain View, CA, USA). Mutation of the DR5 C94 was produced in the DR5 C81S construct to produce the DR5 C81S/C94S using QuikChange mutagenesis and the following primers: 5′-CATATCTCAGAAGACGGTAGAGATAGCATCTCCTGCAAATATGGACAGG-3′ and 5′- CCTGTCCATATTTGCAGGAGATGCTATCTCTACCGTCTTCTGAGATATG-3′. Mutation of the DR5 C137S was introduced into the DR5 C119S construct to produce the DR5 C119S/C137S using QuikChange mutagenesis and the following primers: 5′-CCACGACCAGAAACACAGTGAGTCAGTGCGAAGAAGGCACCTTC -3′ and 5′- GAAGGTGCCTTCTTCGCACTGACTCACTGTGTTTCTGGTCGTGG -3′. The C178S mutation of DR5 was introduced into the DR5 C160S construct to produce the DR5 C160S/C178S using QuikChange mutagenesis and the following primers: 5- CACCCTGGAGTGACATCGAAAGTGTCCACAAAGAATCAGGTAC -3′ and 5′- GTACCTGATTCTTTGTGGACACTTTCGATGTCACTCCAGGGTG-3′. The C153S and C156S mutations of DR5 were produced using QuikChange mutagenesis and the following primers, respectively: 5′-GAAGAAGATTCTCCTGAGATGAGCCGGAAGTGCCGCACAGGG-3′ and 5′- CCCTGTGCGGCACTTCCGGCTCATCTCAGGAGAATCTTCTTC-3′ and 5′- CTCCTGAGATGTGCCGGAAGAGCCGCACAGGGTGTCCCAGAGGG-3′ and 5′- CCCTCTGGGACACCCTGTGCGGCTCTTCCGGCACATCTCAGGAG-3′. The K245R DR5 mutation was produced by QuikChange Mutagenesis and the following primers: 5′- GTCCTTCCTTACCTGCGAGGCATCTGCTCAGGT-3′ and 5′-ACCTGAGCAGATGCCTCGCAGGTAAGGAAGGAC-3′. Tet-DR5 [ΔC41] was produced by amplifying DR5 by PCR, adding 5′-EcoRI and 3′-BamHI sites using the following primers: 5′- TTTTGAATTCCACCATGGAACAACGGGGACAGAAC-3′ and 5′- TTTTGGATCCTTACTTGTCGTCATCGTCTTTGTAGTCGACAGAGGCATCTCGCCCGG-3′ followed by cloning into the EcoRI and BamHI sites of the pRetroX-TetOne-Puro vector. The Tet-DR4[ΔC43] was produced by amplifying DR4 by PCR, adding 5′-ClaI and 3′-BamHI sites using the following primers: 5′-TTTTATCGATCACCATGGCGCCCGTCGCCGTCTGG-3′ and 5′- TTTTGGATCCTTACTTGTCGTCATCGTCTTTGTAGTCGATCGAGGCGTTCCGTCCAGTTTTG-3′ followed by cloning into the ClaI and BamHI sites of the pRetroX-TetOne-Puro vector.

In order to clone ERp44, total RNA from T47D cells was extracted with TRIzol Reagent (Invitrogen, Waltham, MA USA) according to the manufacturer’s protocol. Total cellular RNA was reverse transcribed to synthesize first-strand cDNA using the PCR conditions listed: 25 °C for 10 min, 42 °C for 30 min, and 95 °C for 5 min. DNA encoding ERp44 was subsequently amplified using the following primers: 5′- TTTTGGATCCCACCATGCATCCTGCCGTCTTCC-3′ and 5′- TTTTCTCGAGTTAAAGCTCATCTCGATCCCTC-3′. The PCR fragment encoding ERp44 was cloned into the 5′ BamHI and 3′ XhoI sites of the pMXs-IRES-Blasticidin retroviral vector (RTV-016) (Cell Biolabs, Inc., San Diego, CA USA). The following primers were used to produce the C29S mutation of ERp44 using QuikChange mutagenesis: 5′- CTTTAGTAAATTTTTATGCTGACTGGAGTCGTTTCAGTCAGATGTTG-3′ and 5′- CAACATCTGACTGAAACGACTCCAGTCAGCATAAAAATTTACTAAAG-3.′ All mutations were verified by sequencing.

### MTT Cell Viability Assays

In order to evaluate cell viability, cells were plated at 7,500/well in 96-well plates and incubated at 37°C for 24h. Cells were subsequently treated with various compounds for 72 h at 37°C. Following removal of the cell media, cells were incubated with 0.5 mg/ml MTT (3-(4,5-dimethylthiazol-2-yl)-2,5-diphenyltetrazolium bromide) (Biomatik, Wilmington, DE, United States) in PBS at 37°C for 1 h. The MTT solution was subsequently removed and the MTT formazan product was dissolved in 100 μl of DMSO, followed by measurement of MTT formazan absorbance (570 and 690 nm) in a plate reader.

### Protein Synthesis Assays

Leucine incorporation into proteins was assayed using ^3^H-Leucine (cat. # NET460001MC) obtained from Perkin Elmer (Waltham, MA), as described in a previous publication [33].

### Chemical Synthesis of DDAs

The DDAs presented in Fig. 1A. were prepared based on existing literature procedures from our team and others. RBF3, D5DO, and D7DO were obtained according to the methods described by Field and colleagues [59, 60]. DTDO was synthesized as we described previously [16], as well as tcyDTDO [9], dMtcyDTDO and dFtcyDTDO [61], and Bio-Pyr-DTDO [7].

### Metabolic stability using rat and human liver and intestinal microsomes

To understand the rate of metabolism of compounds across species the *in vitro* metabolic stability of each compound was performed using liver and intestinal microsomes from rats and humans in triplicate. Verapamil was used as a positive control to check the activity of the microsomes. The incubation mixtures consisted of liver or intestinal microsomes (1 mg /ml protein for liver microsomes and 0.5 mg/ml protein for intestinal microsomes), substrate (10 μM), and NADPH (1 mM) in a total volume of 0.2 ml potassium phosphate buffer (50 mM, pH 7.4). Reactions were initiated with the addition of NADPH and kept in an incubator shaker at 37°C. Aliquots of 20 μl were collected at 0, 5, 10, 15, 30, 45, and 60 min and mixed with 100 μl of acetonitrile with formic acid (0.1% v/v) containing phenacetin (50 ng/ml; internal standard) for the termination of the reaction. The samples were then vortex mixed and filtered through a 0.45 μm PTFE Solvinert membrane filtration plate under centrifugation at a speed of 2000 ×g for 5 min at 4°C. The filtrates were subjected to UPLC-MS/MS analysis.

The intrinsic clearance of the compounds was calculated using a half-life employing the ‘substrate depletion’ approach. The apparent half-life was calculated from the pseudo-first-order rate constants obtained by linear regression of log (concentration) and time plots. The *in vitro* intrinsic clearance for compound was estimated using the formula:

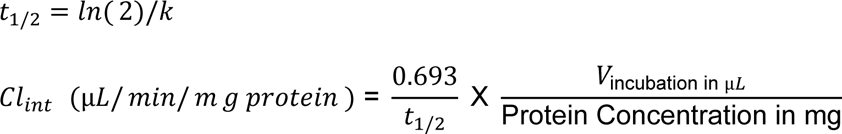

Where k is the slope of the line obtained by plotting the natural logarithmic of the percentage of parent remaining versus time and V is the volume of incubation.

The *in vitro* intrinsic clearance from rat and human liver microsomes were scaled to whole-organ (hepatic) *in vivo* intrinsic CL (CL_int, H_) using the scaling factors available in the literature using equation [62]:

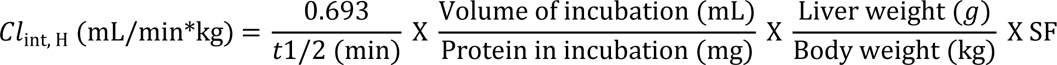

The scaling factor used for the rat was 45 (45 mg microsomal protein/g liver) and liver weight (g) per kg body weight was 40 g/kg while for human scaling factors was 29 (29 mg microsomal protein/g liver) and liver weight (g) per kg body weight was 24 g/kg [63, 64].

### LC-MS/MS analysis

UPLC-MS/MS analysis was carried out using a Waters Acquity Class I Plus UPLC coupled with a Waters Xevo TQ-S Micro triple quadrupole mass spectrometer. The chromatographic separation was achieved using Acquity UPLC CSH C18 column (2.1 mm x 50 mm, 1.7 µm) using the mobile phase consisting of 0.1% formic acid (A) – methanol (B) with a gradient program of 80 % A held for 0.5 min, then decreased A to 65% reaching 1.0 min and further decreased to 40 % A by 2.5 min and held at 40 % until 3.0 min, then sharply decreased back to the initial conditions by 3.1 min and maintained until 3.5 min. The column and autosampler temperatures were kept at 50 °C and 4 °C, respectively. The mobile phase was delivered at a flow rate of 0.35 mL/min and the injection volume was set to 2 μL. The MassLnyx software version 4.2 was used for instrument control and TargetLynx for data analysis. The mass spectrometer was operated in positive ion mode and detection of the ions was performed in the multiple reaction monitoring (MRM) mode. The monitored ion transitions (*m/z*) and instrument conditions can be seen in Table 1. Each compound was monitored using two precursor-to-daughter ion transition pairs, one as a quantifier and another as a qualifier to get better selectivity for each compound. The ion spray voltage was set at 3000 V, the desolvation temperature was 400 °C, the desolvation gas flow was 850 L/h, and the cone gas flow was 50 L/h.

**Table 1.** Mass parameters for tcyDTDO, dMtcyDTDO, dFtcyDTDO, and internal standard (IS)

| Compound | Parent<br>(*m/z*) | Daughter<br>(*m/z*) | Cone<br>(V) | Collision<br>(V) | Type |
| --- | --- | --- | --- | --- | --- |
| tcyDTDO | 207.16 | 81.03 | 46 | 24 | Qualifier |
| tcyDTDO | 207.16 | 109.02 | 46 | 14 | Quantifier |
| dFtcyDTDO | 243.10 | 105.00 | 44 | 20 | Qualifier |
| dFtcyDTDO | 243.10 | 125.10 | 44 | 16 | Quantifier |
| dMtcyDTDO | 267.10 | 139.10 | 34 | 11 | Qualifier |
| dMtcyDTDO | 267.10 | 235.00 | 34 | 4 | Quantifier |
| Phenacetin | 180.11 | 110.02 | 34 | 20 | IS |

### Flow Cytometry Analysis

Cells were lifted from plates using cell scrapers and washed in ice cold PBS. Single cell suspensions were prepared, counted, and diluted to 1 × 10^6^ cells/100 μL. Subsequently, cells were stained for DR4 (DJR1-APC, Cat: 307208, Biolegend) and DR5 (DJR2-4-PE, Cat:307406, Biolegend) markers for 30 min at 4 °C. Cells were then washed twice in ice-cold PBS and stained with viability dye (violet fluorescent reactive dye, Cat:L34955, Invitrogen). FACS Buffer (1% FBS, 0.5 mM EDTA in PBS (400 μL)) was subsequently added. Cells were not fixed or permeabilized. Stained samples were analyzed using single-color compensation and FMO controls on a Sony SP6800 spectral analyzer and quantified using FlowJo V10.8.1 (BD Biosciences). Cells were gated in the following sequence: SSC-A x FSC-A, FSC-H x FSC-A, SSC-H x SSC-A, and Live Cells, to determine Mean Fluorescence Intensity (MFI) of DR5 or DR4.

### Statistical Analysis

Statistical analysis of protein levels detected by immunoblot, MTT viability assays and protein synthesis assays were performed as described in a previous publication [33].

## Acknowledgements

This work was funded in part by the following grants to BL and RC: NIH/NCI R21 CA252400, NIH/NCI R21 CA277485, Florida Department of Health, James & Esther King Cancer Research Program grants 23K06 and 22K04, Florida Department of Health, Bankhead-Coley Research Program grant 23B03, and a grant from the Florida Breast Cancer Foundation. OG was funded by R01 DK121831. We thank the UF Molecular pathology Core, UF ICBR Proteomics Core, UF CTSI Translational Drug Development Core, and UFHCC Flow Cytometry Core for their contributions to the manuscript.

## Author Contributions

Performed experiments: MS, ZD, SE, GT, GA, MW, HS, BF, SK, BL

Directed research: ML, AS, OG, JH, BT, RC, BL

Provided key reagents and expertise: JH, BT

Wrote and edited manuscript: ML, ZD, GT, CWC, AS, OG, RC, BL

