## Supplemental Material for "DR5 disulfide bonding as a sensor and effector of protein folding stress"

### Table of Contents

**Supplemental Figure S1:** *Neither CDCP1 overexpression, or inhibition of eEF2 kinase influence DDA induction of DR5 upregulation, DR4/5 oligomerization, or Caspase 8 cleavage.* ............... Page 2

**S2:** *Effects of tcyDTDO treatment on total blood cell counts* .................................................... Page 3

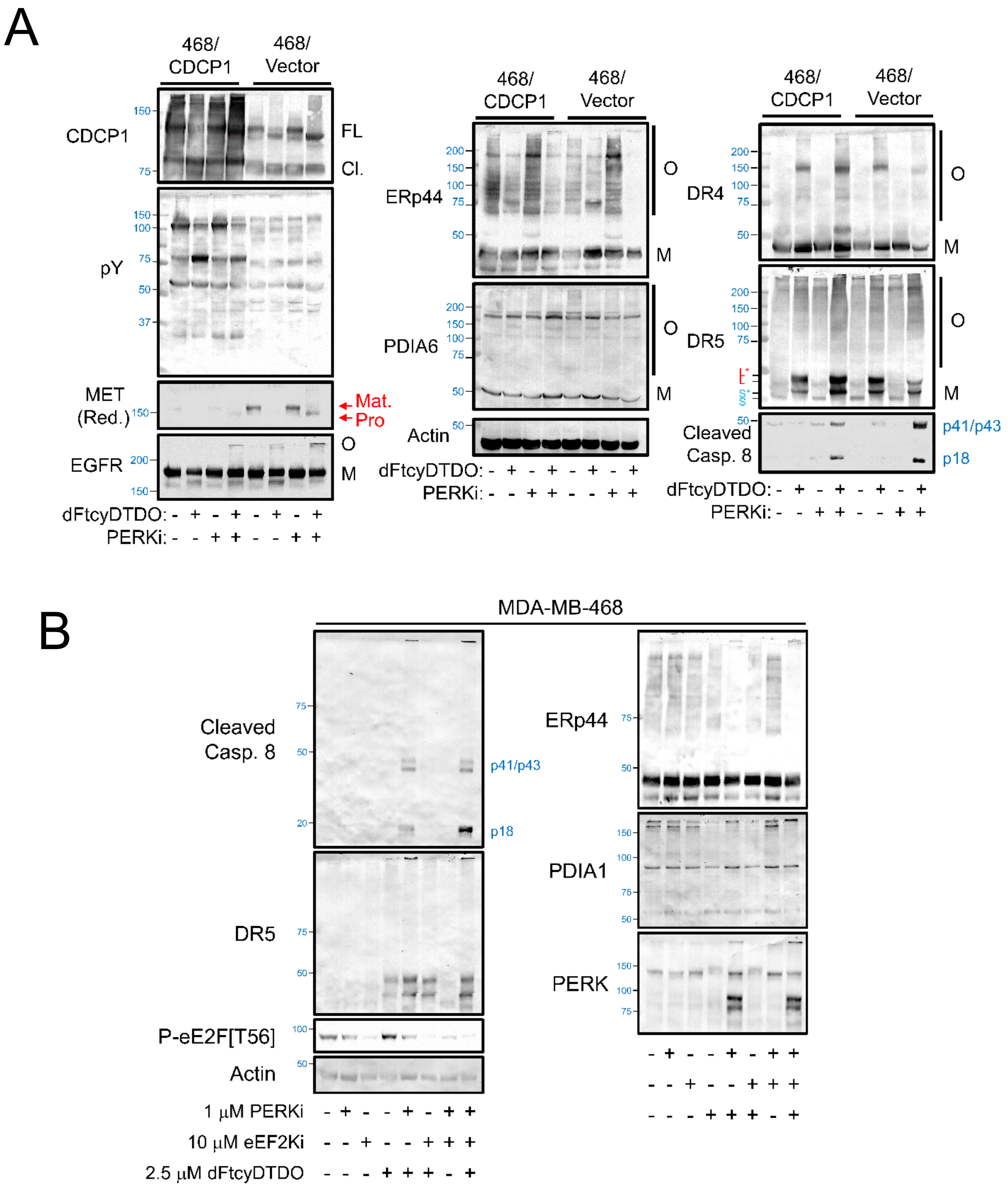

Fig. S1: Neither CDCP1 overexpression, or inhibition of eEF2 kinase influence DDA induction of DR5 upregulation, DR4/5 oligomerization, or Caspase 8 cleavage. MDA-MB-468 cells transduced with CDCP1 vector or empty vector were treated as specified with 2.5 μM dFtcyDTDO or 1 μM PERKI for 24 h and subjected to immunoblot analysis under non-reducing conditions. O and M represent oligomeric and monomeric protein isoforms. FL and Cl. Denote the full length and cleaved forms of CDCP1. B. MDA-MB-468 cells were treated as indicated for 24 h and analyzed by non-reducing immunoblot.

| Sample ID | WBC (10^3/uL) | Neu # (10^3/uL) | Lym # (10^3/uL) | Mon # (10^3/uL) | Eos # (10^3/uL) | Bas # (10^3/uL) | Neu % (%) | Lym % (%) |
|---|---|---|---|---|---|---|---|---|
| VEHICLE 1 | 1.17 | 0.26 | L 0.53 | 0.11 | 0.08 | H 0.19 | 21.9 | 45.1 |
| VEHICLE 2 | 1.35 | 0.29 | 0.84 | 0.08 | 0.06 | 0.08 | 21.1 | 61.8 |
| VEHICLE 3 | 1.8 | 0.28 | 1.37 | L 0.03 | 0.08 | 0.04 | 15.5 | 75.8 |
| tcyDTDO 1 | 10.17 | 0.95 | 7.75 | 0.29 | 0.45 | H 0.73 | 9.3 | 76.2 |
| tcyDTDO 2 | 5.23 | 1.34 | 3.43 | 0.08 | 0.11 | H 0.27 | 25.6 | 65.6 |
| tcyDTDO 3 | 3.13 | 1.04 | 1.67 | 0.1 | 0.21 | 0.11 | 33.2 | 53.3 |
| tcyDTDO 4 | 1.1 | L 0.11 | 0.85 | 0.05 | 0.03 | 0.06 | 9.4 | 77.3 |
| Sample ID | Mon % (%) | Eos % (%) | Bas % (%) | RBC (10^6/uL) | HGB (g/dL) | HCT (%) | MCV (fL) | MCH (pg) |
| VEHICLE 1 | 9.4 | 7.2 | H 16.4 | 8.13 | 12.6 | 37.1 | 45.6 | 15.5 |
| VEHICLE 2 | 6.3 | 4.9 | H 5.9 | 9.06 | 14.1 | 41.5 | 45.9 | 15.6 |
| VEHICLE 3 | 1.8 | 4.3 | H 2.6 | 8.35 | 12.8 | 38.4 | 45.9 | 15.3 |
| tcyDTDO 1 | 2.8 | 4.4 | H 7.3 | 7.65 | 12.2 | 35.6 | 46.5 | 15.9 |
| tcyDTDO 2 | 1.5 | 2.1 | H 5.2 | 9.07 | 13.8 | 41.5 | 45.7 | 15.2 |
| tcyDTDO 3 | 3.2 | 6.6 | H 3.7 | 9.34 | 14 | 42.3 | 45.3 | 15 |
| tcyDTDO 4 | 4.9 | 3.4 | H 5.0 | 6.97 | 11.3 | L 32.6 | 46.7 | 16.2 |
| Sample ID | MCHC (g/dL) | RDW-CV (%) | PLT (10^3/uL) | MPV (fL) | Species | Mode | | |
| VEHICLE 1 | 34 | 12.8 | L 56 | 5.5 | Mouse | Whole Blood | | |
| VEHICLE 2 | 33.9 | 12.8 | L 48 | 5.6 | Mouse | Whole Blood | | |
| VEHICLE 3 | 33.4 | 13.2 | L 48 | 5.8 | Mouse | Whole Blood | | |
| tcyDTDO 1 | 34.3 | 12.8 | L 102 | 5.3 | Mouse | Whole Blood | | |
| tcyDTDO 2 | 33.3 | 12.6 | 684 | 5.4 | Mouse | Whole Blood | | |
| tcyDTDO 3 | 33.1 | 12.3 | L 116 | 5.8 | Mouse | Whole Blood | | |
| tcyDTDO 4 | 34.7 | 12.6 | L 75 | 5.8 | Mouse | Whole Blood | | |

Fig. S2: Total blood cell counts of blood from female mice treated for 20 days with DMSO (Vehicle) and 10 mg/kg dMtcyDTDO). Two hours after the final treatment, , peripheral blood was collected by facial vein puncture into EDTA-treated tubes and complete blood cell counts (CBCs) were obtained. The low platelet numbers likely on both the vehicle and dMtcyDTDO groups likely results from partial clotting of the samples.
